# Efficient long-range conduction in cable bacteria through nickel protein wires

**DOI:** 10.1101/2020.10.23.351973

**Authors:** Henricus T. S. Boschker, Perran L.M. Cook, Lubos Polerecky, Raghavendran Thiruvallur Eachambadi, Helena Lozano, Silvia Hidalgo-Martinez, Dmitry Khalenkow, Valentina Spampinato, Nathalie Claes, Paromita Kundu, Da Wang, Sara Bals, Karina K. Sand, Francesca Cavezza, Tom Hauffman, Jesper Tataru Bjerg, Andre G. Skirtach, Kamila Kochan, Merrilyn McKee, Bayden Wood, Diana Bedolla, Alessandra Gianoncelli, Nicole M.J. Geerlings, Nani Van Gerven, Han Remaut, Jeanine S. Geelhoed, Ruben Millan-Solsona, Laura Fumagalli, Lars-Peter Nielsen, Alexis Franquet, Jean V. Manca, Gabriel Gomila, Filip J. R. Meysman

## Abstract

Filamentous cable bacteria display unrivalled long-range electron transport, generating electrical currents over centimeter distances through a highly ordered network of fibers embedded in their cell envelope. The conductivity of these periplasmic wires is exceptionally high for a biological material, but their chemical structure and underlying electron transport mechanism remain unresolved. Here, we combine high-resolution microscopy, spectroscopy, and chemical imaging on individual cable bacterium filaments to demonstrate that the periplasmic wires consist of a conductive protein core surrounded by an insulating shell layer. The core proteins contain a sulfur-ligated nickel cofactor, and conductivity decreases when nickel is oxidized or selectively removed. The involvement of nickel as the active metal in biological conduction is remarkable, and suggests a hitherto unknown form of electron transport that enables efficient conduction in centimeter-long protein structures.

## Introduction

Cable bacteria are multicellular microorganisms in the Desulfobulbaceae family that display a unique metabolism, in which electrical currents are channeled along a chain of more than 10.000 cells^1–4^. The observation that electrical currents are transported along centimeter-long filaments^3,4^, extends the known length scale of biological electron transport by orders of magnitude, and suggests that biological evolution has resulted in an organic structure that is capable of highly efficient electron transport across centimeter-scale distances^4^. Recent studies demonstrate that cable bacteria effectively harbor an internal electrical grid, which displays a unique topology and exceptional electrical properties^5–8^. The cell envelope of cable bacteria contains a distinctive network of parallel fibers (each ~50 nm diameter) that run along the whole length of the filament^5,6^. These fibers are embedded in a joint periplasmic space and remain continuous across cell-to-cell junctions^5^. Direct electrical measurements demonstrate that these periplasmic fibers are the conductive structures^7,8^. Additionally, the cell-to-cell junctions contain a conspicuous cartwheel structure that electrically interconnects the individual fibers to a central node, and in this way, the electrical network becomes redundant and fail-safe^8^. The electrogenic metabolism of cable bacteria necessitates that nano-ampere currents are efficiently conducted over centimeter scale distances through this network^4^, and in effect, the periplasmic fibers display extraordinary electrical properties for a biological material^7^. The estimated *in vivo* current density of ~10^6^ A m^-2^ is comparable to that of household copper wiring, while the conductivity can exceed 20 S cm^-1^ and thus rivals that of doped synthetic conductive polymers^7^.

Bio-materials typically have an intrinsically low electrical conductivity, and so the availability of a bio-material with extraordinary electrical properties has great potential for new applications in bio-electronics. This prospect of technological application however requires a deeper understanding of the mechanism of electron transport as well as the structure and composition of the conductive fibers in cable bacteria, but at present, these aspects remain highly enigmatic. One important obstacle is that cable bacteria have a highly complex metabolism and life-style, which strongly hampers culturing and biomass collection. The inability to obtain sufficient cable bacteria biomass precludes the implementation of many traditional analytical techniques that could shed light on the chemical composition of the electrical network. Here we solved this problem by the application of high-resolution microscopy, spectroscopy, and chemical imaging methods to individual filaments of cable bacteria. This approach allows to elucidate the chemical structure and composition of the conductive fibers in cable bacteria, and our results reveal that the long-range electron transport in cable bacteria is crucially dependent on proteins containing a sulfur-ligated nickel group. This finding sets the conduction mechanism in cable bacteria apart from any other known form of biological electron transport, and demonstrates that efficient conduction is possible through centimeter-long protein structures.

### Protein fibers on a polysaccharide-rich layer

Through sequential extraction, a so-called fiber sheath can be isolated from the periplasm of cable bacteria filaments^5^, which contains the conductive fibers^7,8^. Previous studies have already elucidated the geometrical configuration of this fiber network^5–8^. Here, we applied High Angle Annular Dark Field - Scanning Transmission Electron Microscopy (HAADF-STEM) and subsequent tomography, which provided additional details of the fiber sheath architecture (Fig. 1A, Supplementary movie S1), demonstrating that the ring of regularly spaced fibers is held together by a basal sheath. The fibers are clearly visible in unstained preparations, thus confirming that they form electron dense structures^2,5^.

**Fig. 1.**
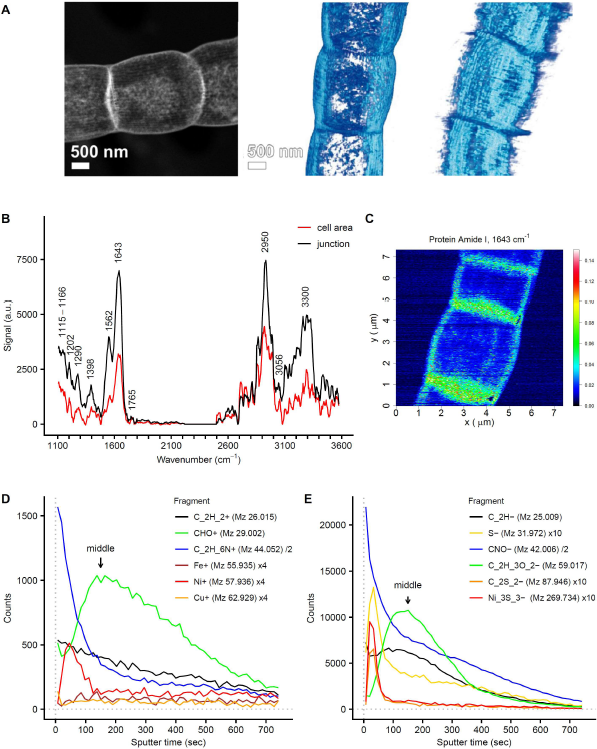
The conductive fiber sheath in cable bacteria is composed of a layer of aromatic-rich protein on top of an acidic polysaccharide layer. A) STEM-HAADF imaging demonstrates that the fiber sheath is composed of parallel fibers imposed on a basal sheath. One 2D image (left panel) and two 3D tomographic reconstructions are shown. B) AFM-IR spectra of fiber sheaths at cell areas and cell junctions (OPO laser, spectra are background corrected and averaged, cell area N = 14, junctions N = 11). C) Fiber sheath AFM-IR mapping of the signal from the 1643 cm^-1^ Amide I protein band (QCL laser, arbitrary units; see Supplementary Figure 7 for corresponding AFM height and deflection images). D-E) Representative ToF-SIMS depth profiles of fiber sheaths obtained in positive (D) and negative mode (E). A selection of fragments from different compound classes is shown (general organic carbon fragments: C_2_H_2_^+^ and C_2_H^-^, protein derived fragments: C_2_H_6_N^+^ and CNO^-^, carbohydrate derived fragments: CHO^+^ and C_2_H_3_O_2_^-^ and sulfur and transition metals). See Supplementary Figure 1 and supplementary materials for further information. Counts of individual fragments were scaled to improve clarity as indicated in the figure legends. The counts from Ni_3_S_3_^-^ are the sum of all ^58^Ni and ^60^Ni isotopologues. Arrows denote the middle of the fiber sheath as calibrated by *in situ* AFM (59 ± 6 nm, see Supplementary Figure 2).

Individual fiber sheaths were subjected to various forms of spectroscopy and chemical imaging. Atomic Force Microscopy - IR spectroscopy (AFM-IR) provided a first insight into the biochemical composition of the fiber sheath (Fig. 1B), and indicated that it mainly consists of protein (N-H/O-H 3300 cm^-1^, Amide I 1643 cm^-1^, Amide II 1562 cm^-1^, Amide III 1290 cm^-1^) and polysaccharide (1202 cm^-1^ and broad feature at 1115-1166 cm^-1^) (band assignment based on^9,10^). The detection of ester groups in the AFM-IR spectra (C=O stretching at 1765 cm^-1^; C=O bending at 1398 cm^-1^) suggested that the polysaccharide contains acidic sugars^9^. Cell junctions give similar AFM-IR spectra as central cell areas, although signals are higher in the junctions (Fig. 1B and 1C,), likely due to presence of the cartwheel structure that interconnects fibers^5^. Mapping of the Amide I band gave a relatively even signal across the central cell area with no indication of a fiber structure (Fig. 1C), thus suggesting that there is protein throughout the fiber sheath. A relatively strong aromatic C-H stretching band at 3056cm^-1^ further suggests that the protein is rich in aromatic amino acids^9,10^, which have been proposed to play a role in the electron transport within *Geobacter* pili^11^. A recent genome and proteome study^12^ also speculates that the periplasmic fibers of cable bacteria could be composed of bundles of pilin protein, as found in the conductive pili of *Geobacter*^11,13^. However, the position of the Amide I peak at 1643 cm^-1^ indicates that the protein secondary structure is mainly disordered, and not of the α-helix type^10^, which hence speaks against an abundance of α-helix rich pili.

Time of Flight-Secondary Ion Mass Spectrometry (ToF-SIMS) analysis was applied in combination with *in situ* AFM calibration of the sputtering depth, and this enabled us to map the nanometer-scale depth distribution of both organic and inorganic constituents within the fiber sheath (additional ToF-SIMS results are provided in the supplementary materials). Replicate ToF-SIMS analyses of fiber sheaths in both positive and negative mode yielded a consistent depth distribution of organic fragments (Fig. 1 D-E, Supplementary Figure 1, Supplementary Table 2 and 3). Initially, high signals were recorded for a variety of amino acid fragments, including all three aromatic amino acids^14,15^. After ~150 sec of sputtering, the amino acid-derived signal levelled off, while counts of oxygen-rich fragments, including carbohydrate specific ions (C_2_H_5_O_2_^+^, C_3_H_3_O_2_^+^ and C_3_H_5_O_2_^+^)^16^ peaked. In these samples, the fiber sheath forms a flattened hollow cylinder upon the supporting substrate, which has a total thickness of 117 ± 10 nm (superposition of top and bottom cell envelope layers as measured in the middle of a cell; Supplementary Figure 2). AFM calibration of the sputtering time places the carbohydrate peak at 59 ± 6 nm depth (Supplementary Figure 2), which matches the middle of the flattened fiber sheath between the top and bottom layers. This suggests that the fiber sheath is made of a protein layer on top of a basal polysaccharide-rich layer.

Together, the HAADF-STEM, AFM-IR and ToF-SIMS data show that (i) the conductive fibers are positioned in a regular, parallel pattern on the outside of the fiber sheath, (ii) that the fibers consist of protein that is rich in aromatic amino acids, and (iii) that the fibers rest upon a basal sheath rich in polysaccharide.

### A sulfur-ligated metal group

To obtain further insight into the composition of the conductive periplasmic fibers, Raman microscopy with different laser wavelengths was applied to both intact cable bacteria and extracted fiber sheaths. Green laser (523 nm) Raman microscopy spectra obtained from living cable bacteria showed the resonance bands of cytochrome heme-groups (750, 1129, 1314 and 1586 cm^-1^) seen previously^3^, but additionally revealed two prominent bands within the low-frequency region at 371 and 492 cm^-1^ (Fig. 2A). These two Raman bands remained prominently present when cable bacterium filaments were isolated from the sediment and air-dried. Moreover, the two bands appeared in both thick (~4 μm diameter) and thin (~1 μm diameter) filament morphotypes, as well as in marine and freshwater cable bacteria (Fig. 2A, Supplementary Figure 3A). This suggests that the two low-frequency bands are a core feature of the cable bacteria clade. The low-frequency position points towards a cofactor that involves a heavy atom, such as a ligated metal group^17^.

**Fig. 2.**
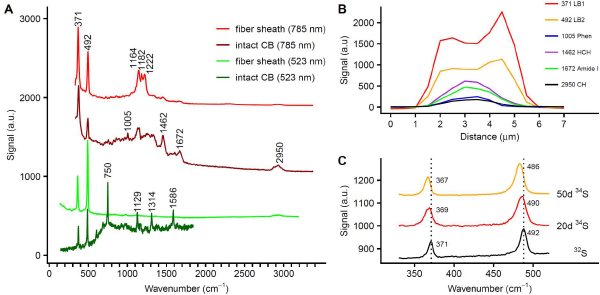
Raman spectra of intact cable bacteria and fiber sheaths indicating a sulfur-ligated metal group in the fiber sheath. A) Raman spectra collected with green (523 nm) and NIR (785 nm) lasers. The low frequency bands at 371 and 492 cm^-1^ indicate the presence of a metal group, and are present in all spectra. The dark green spectrum is from intact, living cable bacteria (CB) in a gradient slide, while the dark red spectrum is recorded on intact, dried cable bacterium filaments. The light green and light red spectra are from fiber sheaths. B) Variation of the Raman signal (NIR laser) in a transversal section across an intact, dried cable bacterium filament, which was ca. 3 μm wide. The most prominent bands are shown: the two low-frequency bands (371 LB1, 492 LB2), phenylalanine ring-breathing (1005 Phen), CH_2-_bending (1462 HCH), the protein Amide I band (1672 Amide I) and CH-stretching (2950 CH). C) Average Raman spectra (green laser) and peak shifts resulting from ^34^S labelling of intact, dried cable bacterium filaments. The two low-frequency bands are shown after 20 and 50 days of incubation and compared to the unlabeled control spectrum.

When air-dried intact cable bacteria were investigated with near infrared (NIR) laser (785 nm) Raman spectroscopy, the two low-frequency bands were again present, but the spectra showed additional bands (Fig. 2A), characteristic of general biomolecular constituents, such as C-H stretching (2950 cm^-1^), the Amide I peak of protein (1672 cm^-1^), CH_2_ bending (1462 cm^-1^), and phenylalanine ring-deformation (1005 cm^-1^)^18^. Cross-sectional scans of intact filaments showed a unimodal signal for biomolecular signals generally present in microbial biomass (phenylalanine, C-H bonds, Amide I). In contrast, the signal of the two low-frequency bands showed a bimodal maximum at the filament edges (Fig. 2B), which suggest that the metal moiety is located in the cell envelope, possibly in the periplasmic fiber sheath (Supplementary Figure 3B). This was confirmed by Raman spectroscopy of fiber sheaths extracted from intact bacteria (Fig. 2A), which produced simple green-laser spectra that only contained the two low-frequency bands and a weak C-H signal at 2950 cm^-1^. Furthermore, these spectra also showed no cytochrome signal^7^, thus confirming that the conduction mechanism does not involve cytochromes as seen in *Shewanella* and some types of *Geobacter* nanowires^19,20^. NIR laser Raman spectra of fiber sheaths additionally revealed three conspicuous bands at 1164, 1182 and 1222 cm^-1^ (Fig. 2A), which may originate from single carbon bonds (C-C, C-O or C-N) associated with the metal group^17^, and also showed small bands from protein (Amide I, 1665 cm^-1^) and C-H (1451 and 2950 cm^-1^)^18^. Ratios between the background-corrected peak heights of the two low-frequency bands were similar in all Raman spectra recorded (green-laser R_492/371_ = ~2.1, NIR-laser R_492/371_ = ~0.6), indicating that both bands originate from a single moiety.

To further examine the origin of the two low-frequency bands, we grew cable bacteria in sediments amended with ^34^S or ^13^C stable isotope tracers and investigated intact air-dried filaments with Raman spectroscopy as before. Labelling with ^34^S did not affect the cytochrome bands as expected, but resulted in a shift in both low-frequency bands towards lower wave numbers (Fig. 2C). This indicates that sulfur is directly involved in both low-frequency bands and the metal group therefore appears to be S-ligated. Labelling with ^13^C resulted in substantial shifts of the cytochrome bands to lower values, as expected. This also demonstrated that cable bacteria were highly labelled (Supplementary Figure 4), but nevertheless, the 371 cm^-1^ band showed no response to ^13^C labelling, while the 492 cm^-1^ band only displayed a small shift to lower wave numbers. This suggests that carbon is not directly involved in in metal ligation, but could be present further away (e.g. by having carbon atoms adjacent to the sulfur-ligated metal group).

### Fibers are enriched in nickel and sulfur

To identify the metal in the sulfur-ligated group, we first analyzed the elemental composition of cable bacterium filaments by STEM-Energy-Dispersive X-ray spectroscopy (EDX) (Fig. 3A-B, Supplementary Table 1). Metals commonly found in metalloproteins were present in intact filaments, but concentrations were low and close to detection limits: Fe (0.033-0.047 Atm%), Ni (0.009 Atm%) and Cu (0.006-0.009 Atm%). After fiber sheath extraction, Ni (0.016-0.037 Atm%) was selectively enriched by a factor of 2-4 compared to intact filaments. In contrast, Fe was partially removed by a factor of ~2, consistent with the loss of cytochromes, while Cu remained equally low. Additional metal analysis by Synchrotron Low-Energy X-Ray Fluorescence (LEXRF) (Fig. 3C-E) showed that absolute Ni counts were similar in intact bacteria and fiber sheaths, thus confirming that Ni is concentrated in the fiber sheath. Fe was again selectively lost during extraction of fiber sheaths, while Cu levels were highly variable and exceeded STEM-EDX values, suggesting that Cu data were affected by contamination during LEXRF analysis.

**Fig. 3.**
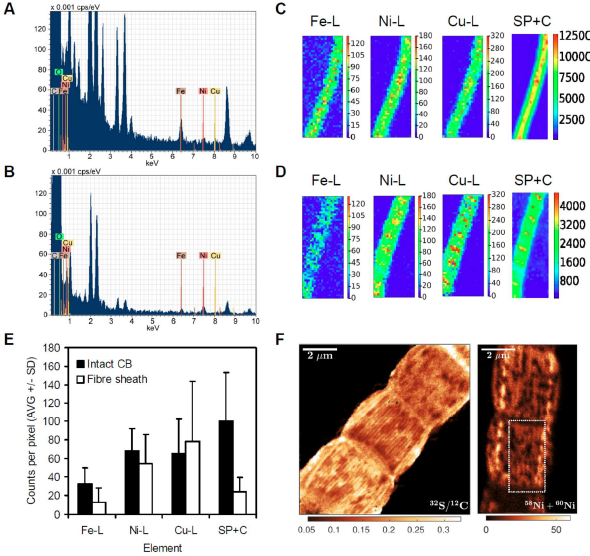
Elemental analysis shows that the fibers are Ni and S rich. Representative STEM-EDX spectra from A) intact cable bacteria and B) fiber sheaths shows a detectable Ni signal and lower Fe and Cu levels in the fiber sheath. Elemental compositions are found in Supplementary Table 1. Representative synchrotron LEXRF maps for C) intact cable bacteria (10 μm x 25 μm) and D) fiber sheaths (11 μm x 27 μm). SP+C denotes Scatter Peak plus Compton and L denotes low-energy L-band. E) Average counts per pixel from LEXRF maps showing that Ni is mainly found in the fiber sheath (intact cable bacteria N = 5 and fiber sheaths N = 6, background corrected). Data given for the detected transition metals and SP+C. The latter data were scaled to fit into the graph by setting the average of the intact cable bacteria (CB) counts to 100 (original counts 4290 ± 2640). F) Nano-SIMS images of fiber sheaths. Mapping of ^32^S/^12^C ion count ratio (first 100 planes) shows the sulfur rich fibers. The Ni (^58^Ni+^60^Ni) ion count has a lower signal/noise ratio and its mapping (first 50 planes) only shows visible fibers is restricted regions (as indicated by the rectangle). The complete set of Nano-SIMS images is given in Supplementary Figure 5.

Together, the STEM-EDX and LEXRF data indicated that Ni was the most likely candidate for the metal contained in the sulfur-ligated group. This hypothesis was confirmed by ToF-SIMS analysis of the fiber sheaths (Fig. 1D, a detailed discussion of ToF-SIMS data is given in supplementary text). In positive mode the four main Ni isotopes (^58^Ni, ^60^Ni, ^61^Ni and ^62^Ni) showed a sharp subsurface peak, while the minor isotope ^64^Ni showed mass interference (likely from a low amount of ^64^Zn). Other transition metals had either low counts (Cu and Fe, Fig. 1D) or were not detectable (Mn, Co and Mo). Negative mode ToF-SIMS depth profiles showed a subsurface peak of various S-derived anions (^32^S^-^, ^34^S^-^, SH^-^ and S_2_^-^) at the same position as the Ni peak (Fig. 1E), in agreement with a sulfur-ligated Ni group. The Ni and S peak emerged after 33-46 sec of sputtering within the first fiber protein layer (corresponding to 15 ± 3 nm of sputtering depth, Supplementary Figure 2), and was preceded by thin proteinaceous surface layer devoid of Ni.

High resolution Nano-SIMS analysis confirmed that that the conductive fibers were Ni and S rich. S^-^-ion maps revealed a parallel line pattern (Fig. 3F) with a similar line spacing (~150 nm) between fibers as reported previously (150-200 nm)^5^. Although metal maps generally have a lower signal-to-noise ratio, the ^58^Ni^+^+^60^Ni^+^ signal did show an indicative line spacing of ~200 nm in restricted areas (Fig. 3F). Furthermore, the presence of NiS-cluster ions such as Ni_3_S_3_^-^ in the negative mode ToF-SIMS spectra (Fig. 1E, Supplementary Figure 9) also suggests that Ni and S must be present in close proximity (lateral distance (XY) within <0.5 nm, depth (Z) within <2 nm), otherwise these NiS-clusters would not form in the ToF-SIMS ion plume^21^. Finally, we detected two organic sulfur fragments (C_2_S_2_^-^ and C_2_S_2_H^-^) that were specifically associated with the Ni_3_S_3_^-^ peak (Fig. 1E, Supplementary Figure 1). This could indicate that the fibers are rich in proteins with disulfide bonds, which may in part explain their high sulfur content and their chemical resistance (i.e. the fibers survive the SDS/EDTA extraction procedure). Alternatively, these two organic fragments could have come from the Ni ligating group. Combined, our results demonstrate that the individual fibers are Ni and S rich and that Ni represents the metal in the sulfur-ligated group as detected by Raman analysis.

### The Ni/S group and long-distance electron transport

To verify whether the Ni/S-group truly plays a role in long-distance electron transport, we first studied the effect of redox state on Raman signals and conductance. Chemically reducing the fiber sheath with K_4_Fe^II^(CN)_6_ resulted in a small increase in the green laser Raman signal of the two low-frequency bands, while oxidizing the fiber sheath with K_3_Fe^III^(CN)_6_ almost completely removed these signals (Fig. 4A). Such oxidation state dependent Raman behavior is commonly observed in metalloproteins, such as cytochromes and [FeNi]-hydrogenases, where only one state shows a high resonance Raman signal^3,22^. Subsequent reduction with K_4_Fe^II^(CN)_6_ restored the Raman signal, suggesting that the Ni/S group is a reversible redox group. Intriguingly, the conductance of the fiber sheath was also reversibly affected by the redox state of the Ni/S group (Fig. 4B). Reduced fiber sheaths showed a 2.1 ± 0.5 (N = 11) higher conductance than oxidized fiber sheaths (independent of the direction of the oxidation/reduction step). The decrease of conductance upon oxidation is consistent with previous observations that the conductance of the fiber sheath decreases in ambient air^7^, suggesting that the Ni/S group is oxidized upon exposure to oxygen, inducing a loss of conductivity.

**Fig. 4.**
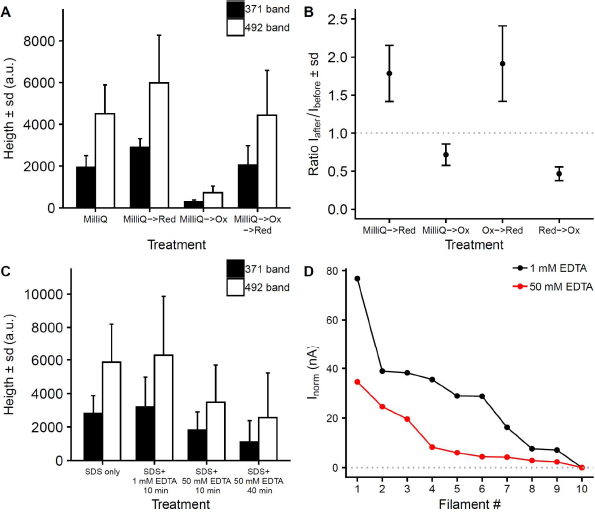
Redox and Ni-removal experiments indicate that the Ni/S group plays a role in electron conduction. A) The effect of oxidation and reduction on green-laser Raman signals from the sulfur-ligated Ni group. The MilliQ->Red treatment was significantly higher than the MilliQ treatment (p < 0.05) and the MilliQ->Ox treatment was significantly lower than all other treatments (p < 0.001, Wilcoxon test, N = 24 to 33). The ratio between the 371 and 492 cm^-1^ Raman bands was not affected by oxidation or reduction. B) The effect of oxidation and reduction treatments on the conductance of individual fiber sheaths. The ratio of the electrical current (I) through the fiber sheath is plotted before and after treatment. The effect in all treatment pairs was significant (p<0.01, Wilcoxon test, N = 4 to 6, tested against no effect Ratio = 1). C) The effect of EDTA with on green-laser Raman signals from the sulfur-ligated Ni group. The decrease in Raman signal between the standard protocol and both high EDTA treatments was significant (p<0.05, Wilcoxon test, N = 14 to 19). D) The effect of high EDTA extraction on normalized conduction of individual fiber sheaths (I_norm_: electrical current normalized to filament length 0.3 mm and bias 0.1 V). Fibers sheaths were extracted with the standard protocol (1% SDS + 1 mM EDTA 10 min) and high EDTA treatment (1% SDS + 50 mM EDTA 10 min). The decrease in I_norm_ between the standard protocol and the high EDTA treatment was significant (Wilcoxon test, p = 0.041, N = 10).

Additional experiments, in which Ni was partially removed from the fiber sheath through extraction with high EDTA concentrations, confirmed that the Ni/S-group plays a crucial role in electron transport. Extraction with 50 mM EDTA left the fiber structure intact (Supplementary Figure 6), but decreased the Raman signal by 45%, indicating that Ni was selectively removed (Fig. 4C), and concomitantly reduced the conduction by 62% (Fig. 4D). This confirms that the Ni/S-group plays a key role in maintaining high rates of long-distance electron transport in cable bacteria.

### The core-shell model of a conductive fiber

By combining and integrating the various types of compositional data collected, we can construct a chemical model of the conductive fiber sheaths in cable bacteria (Fig. 5). The fibers are found on the outside of the fiber sheath and primarily consist of protein (Fig. 1). On the cytoplasmic side of the fiber sheath, the fibers are embedded in or attached to a polysaccharide-rich layer (Fig. 1), most likely made of peptidoglycan as commonly found in Gram-negative bacteria^23^ (see supplementary text for further discussion). This polysaccharide layer holds the fibers together and possibly adds tensile strength to the fiber sheath, which can withstand high pulling forces during filament extraction. ToF-SIMS analysis (Fig. 1D and E) suggests that the fibers themselves are composed of two distinct regions. The central core of the fiber contains protein material that is rich in Ni, while it is also surrounded by a thin layer of Ni deficient protein (Fig. 5A). This core/shell model is consistent with recent conductive AFM investigations of fiber sheaths, which reveal that fibers only display electrical conductivity when a non-conductive surface layer is first etched away^8^. The cross-sectional structure of the fibers therefore resembles a standard household electrical wire, with a conductive core surrounded by electrically insulating layer. We speculate that this insulating layer prohibits that electrons go astray during long-range transport, thus avoiding radical formation and damage to the surrounding cell environment.

**Fig. 5.**
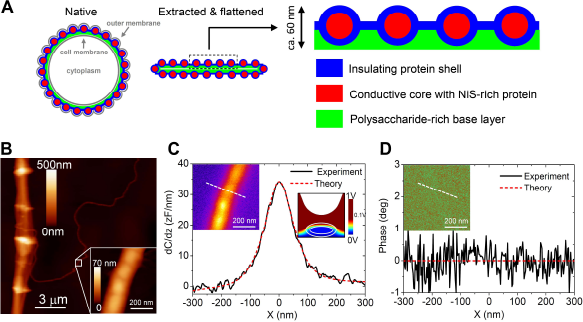
Fiber sheath model and electrostatic properties. A) Compositional model of the conductive fiber sheath in cable bacteria based on the present findings. Cross-sections through a filament in the middle of a cell are drawn and the number of fibers has been reduced for clarity - a 4 μm diameter cable bacterium has typically ~60 fibers^5^. In its native state (right panel), the fiber sheath is embedded periplasm between the cell and outer membrane and adopts a circular shape. After extraction, which removes the membranes and most of the cytoplasm and after drying upon a surface for analysis, the fiber sheath flattens, leading to two mirrored sheaths on top of each other (middle panel). The enlargement shows a section of the top sheath, which is the sample section probed by ToF-SIMS depth profiles and NanoSIMS images. Fibers are made of protein with a conductive Ni/S rich core and a non-conductive outer shell, and are embedded in a basal layer enriched in polysaccharide. B) Topographic AFM image of a fiber sheath with a single isolated fiber detaching. The insert shows a detailed AFM image of this single fiber. C) SDM amplitude image (right insert) and cross-sectional profile. D) Corresponding SDM phase image (insert) and cross-sectional profile. Constant height (z=66 nm) cross-section profiles are measured along the dashed lines shown in the left inserts. The red dotted lines in C) and D) represent model fits assuming the a fiber has a conductive core and an insulating outer shell. The right insert in panel C shows a vertical cross-section of the electric potential distribution as predicted by the model. Model parameters: shell thickness, d=12 nm; fiber height, h=42 nm; fiber width w=87 nm; relative dielectric constants of the shell and core, ε_s_= ε_c_=3; conductivity of the shell σ_s_=0 S/cm (insulating); conductivity of the core σ_c_=20 S/cm ^7^ (see supplementary text for treatment of SDM results and models tested).

Our fiber core/shell model was independently verified by Scanning Dielectric Microscopy (SDM), which enables AFM-based electrostatic force detection^24,25^. We analyzed single, isolated fibers that had separated from a fiber sheath (Fig. 5B), and interpreted the modulus and phase (Figs. 5C and 5D) of the 2ω-electric force harmonic with a computational finite-element model of a flattened cylindrical fiber (right insert in Fig. 5C; height = 41 nm; width = 87 nm, obtained from the deconvoluted topographic image in the insert of Fig. 5B). When this model assumed that the fiber was conductive (σ_c_ = 20 S cm^-1^ as determined in ^7^) and homogeneous (only core, no shell), it could not fit both the modulus and phase data of the electric force (see supplementary text and Supplementary Figure 13 for details). Alternatively, when we assumed the fiber was homogenous and non-conductive (σ_c_ = 0 S cm^-1^), this resulted in an anomalously high relative permittivity ε_r_ = 11 ± 3, implying that the dielectric response of the fiber material would substantially exceed the typical values for common proteins (ε_r_ = 3-5)^24–27^, and would even surpass that of nucleic acids (ε_r_ = ~8)^24,25^. This is not congruent with our AFM-IR and Tof-SIMS data, which demonstrate that the fibers are made of protein. However, when we parameterized the fiber model to include a conductive protein core (ε_c_ = 3; σ_c_ = 20 S cm^-1^) surrounded by a non-conductive protein shell (ε_s_ = 3; conductivity σ_s_ = 0 S cm^-1^), we could fit both the modulus and phase data of the electric force, arriving at a shell thickness d=12±2 nm (red dashed lines in Figs. 5C and 5D). The SDM data therefore add further support to the proposed core/shell model (see supplementary text for a full description of SDM results and models tested).

## Discussion

Our results demonstrate that conduction in cable bacteria occurs through proteins with Ni-dependent cofactors. The observation that Ni plays a crucial role in long-range biological conduction is remarkable, as biological electron transport typically involves Fe and Cu metalloproteins^28^, though not enzymes with Ni-centers. Nickel acts as a catalytic center in only nine enzymes, which are mostly involved in the metabolism of gases^29–31^, but not in electron transport. Clearly, we are dealing with a new type of Ni cofactor, as the S-ligated Ni-group in the periplasmic fibers has a well-defined Raman signature, which does not resemble that of any of the known sulfur-ligated nickel enzymes^22,32^.

Our data also provide a first insight into the structure of this Ni-dependent cofactor. The low-frequency band at 371 cm^-1^ (Fig. 2) is most likely due to Ni-S bond stretching, and bands at similar wave numbers are found in S-ligated Ni-metalloproteins^22,32^ and NiS minerals^33–35^. However, these spectra typically show additional smaller bands from other Ni-S vibrational modes^22,32–35^, which are absent in the fiber sheath spectra (Fig. 2). Our stable isotope labelling data show that the second low-frequency band at 492 cm^-1^ also must involve sulfur and maybe indirectly carbon. While a similar band is observed in NiS_2_ mineral spectra, these also show additional bands around 270 cm^-1^ ^33,35^ that are not seen in the fiber sheath spectra, leaving the alternative possibility that the 492 band derives from S-S stretching^17^. Finally, both the green and NIR Raman spectra show some resemblance to Ni-bis-dithiolene ligands^36,37^. The 492 band then would come from ring-breathing of the aromatic ring containing the dithiolene, and the middle bands in the NIR Raman spectrum (at 1164, 1182 and 1222 cm^-1^, Fig. 2A) could originate from C-C or C-N stretching in the aromatic ring^36,37^. Also, the two organic sulfur fragments associated with the conductive core as detected by ToF-SIMS could possibly be derived from a dithiolene ligand (C_2_S_2_^+^ could be S-C=C-S^+^, for instance). Future studies should better resolve the Ni center coordination of this novel cofactor, and clarify its role in electron transport.

While protein is generally considered to be an electrical insulator, recent work demonstrates that electrical currents can propagate efficiently through nanometer-thick protein films sandwiched between electrodes^38,39^, as well as through micrometer-scale appendages of metal-reducing bacteria that either consist of pilin- or cytochrome-based material^19,40,41^. Our results now extend this known length scale of protein conduction from micrometers to centimeters. At this moment, the exact mechanism of conduction remains unclear, but our results demonstrate that the novel Ni-cofactor is an essential component. Moreover, as we also detected substantial signals from aromatic amino acids in the fiber proteins, one possibility is that conduction is based on electron transfer between S-ligated Ni-groups assisted by bridging aromatic groups in nearby aromatic amino acids.

Together, our data suggest that highly efficient conduction in cable bacteria talks place through proteins with Ni-dependent cofactors, thus providing a mechanism of long-range electron transport that is hitherto unknown to science. This observation that cable bacteria can naturally assemble long, lightweight, flexible, and strong protein wires with exceptional electrical properties potentially opens a promising gateway for new technology, and creates the prospect of bio-electronic devices with new functionality that integrate proteins as new class of electronic materials.

## Acknowledgements

The authors thank Marlies Neiemeisland for assistance with Raman microscopy, Michiel Kienhuis for assistance with NanoSIMS analysis, Peter Hildebrandt and Diego Millo for helping with the interpretation of the Raman spectra, ION-TOF for the Orbitrap Hybrid-SIMS analysis, and Rene Fabregas for helping with finite-element numerical modelling for SDM.

## Funding

HTSB and FJRM were financially supported by the Netherlands Organization for Scientific Research (VICI grant 016.VICI.170.072). Research Foundation Flanders supported FJRM, JVM and RTE through FWO grant G031416N, and FJRM and JSG through FWO grant G038819N. NMJG is the recipient of a Ph.D. scholarship for teachers from NWO in the Netherlands (grant 023.005.049). The NanoSIMS facility at Utrecht University was financed through a large infrastructure grant by the Netherlands Organization for Scientific Research (NWO, grant no. 175.010.2009.011) and through a Research Infrastructure Fund by the Utrecht University Board. AGS is supported by the Special Research Fund (BOF) of Ghent University (BOF14/IOP/003, BAS094-18, 01IO3618) and FWO (G043219). The ToF-SIMS was funded by FWO Hercules grant (ZW/13/07) to JVM and AF. HL, RMS and GG were funded by the European Union H2020 Framework Programme (MSCA-ITN-2016) under grant agreement n°721874.EU, the Spanish Agencia Estatal de Investigación and EU FEDER under grant agreements TEC2016-79156-P and TEC2015-72751-EXP, the Generalitat de Catalunya through 2017-SGR1079 grant and CERCA Program. GG was recipient of an ICREA Academia Award, and HL of a FPI fellowship (BES-2015-074799) from the Agencia Estatal de Investigación/Fondo Social Europeo. LF received funding from the European Research Council (grant agreement No. 819417) under the European Union’s Horizon 2020 research and innovation programme.

## Author contributions

HTSB and FJRM designed the study and were involved in all experiments, measurements and data analysis. SHM performed filament cultivation and fiber sheath extraction. SHM, HTSB, NVG and HR were involved in the Ni group redox and high EDTA experiments. PLMC, KK and BW performed the stable isotope labelling experiments. LP and NMJG performed the NanoSIMS analysis. HTSB, RTE, VS, AF and JVM carried out the ToF-SIMS analysis. HTSB, PLMC, KK, BW, DK, JTB and AGS performed the Raman analysis. NC, PK, DW and SB carried out the HAADF-STEM and STEM-EDX analysis. KKS, FC and TH performed the AFM-IR analysis. DB, AG and SHM carried out the LEXRF measurements. HTSB and FJRM developed the chemical model of the fiber sheath with additional input from JSG, NVG and HR. HL and RMS carried out the SDM experiments and analyzed the results. LF and GG developed the SDM methods and supervised the interpretation of the SDM results. HTSB and FJRM wrote the paper with contributions provided by all co-authors.

## Competing interest declaration

The authors declare that they have no conflict of interest.

## Supplementary information

Methods

Supplementary text

Supplementary Fig. 1 to 16

Supplementary Tables 1 to 3

Supplementary movie: Movie S1.mp4

## Notes

### Competing Interest Statement

The authors have declared no competing interest.

## Reference

1. Nielsen, L. P., Risgaard-Petersen, N., Fossing, H., Christensen, P. B. & Sayama, M. Electric currents couple spatially separated biogeochemical processes in marine sediment. Nature 463, 1071–1074 (2010).

2. Pfeffer, C. et al. Filamentous bacteria transport electrons over centimetre distances. Nature 491, 218–221 (2012).

3. Bjerg, J. T. et al. Long-distance electron transport in individual, living cable bacteria. Proc. Natl. Acad. Sci. 115, 5786–5791 (2018).

4. Meysman, F. J. R. Cable bacteria take a new breath using long-distance electricity. Trends Microbiol. (2018) doi:10.1016/j.tim.2017.10.011.

5. Cornelissen, R. et al. The cell envelope structure of cable bacteria. Front. Microbiol. 9, (2018).

6. Jiang, Z. et al. In vitro single-cell dissection revealing the interior structure of cable bacteria. Proc. Natl. Acad. Sci. 201807562 (2018) doi:10.1073/pnas.1807562115.

7. Meysman, F. J. R. et al. A highly conductive fibre network enables centimetre-scale electron transport in multicellular cable bacteria. Nat. Commun. 10, 1–8 (2019).

8. Eachambadi, R. T. et al. An Ordered and Fail-Safe Electrical Network in Cable Bacteria. Adv. Biosyst. 2000006 (2020) doi:10.1002/adbi.202000006.

9. Naumann, D. Infrared spectroscopy in microbiology. Encycl. Anal. Chem. (2000).

10. Barth, A. Infrared spectroscopy of proteins. Biochim. Biophys. Acta BBA - Bioenerg. 1767, 1073–1101 (2007).

11. Malvankar, N. S. et al. Structural basis for metallic-like conductivity in microbial nanowires. mBio 6, e00084–15 (2015).

12. Kjeldsen, K. U. et al. On the evolution and physiology of cable bacteria. Proc. Natl. Acad. Sci. 201903514 (2019) doi:10.1073/pnas.1903514116.

13. Lampa-Pastirk, S. et al. Thermally activated charge transport in microbial protein nanowires. Sci. Rep. 6, 23517 (2016).

14. Baugh, L. et al. Probing the orientation of surface-immobilized protein G B1 using ToF-SIMS, sum frequency generation, and NEXAFS spectroscopy. Langmuir 26, 16434–16441 (2010).

15. Lebec, V., Boujday, S., Poleunis, C., Pradier, C.-M. & Delcorte, A. Time-of-flight secondary ion mass spectrometry investigation of the orientation of adsorbed antibodies on SAMs correlated to biorecognition tests. J. Phys. Chem. C 118, 2085–2092 (2014).

16. Goacher, R. E., Jeremic, D. & Master, E. R. Expanding the library of secondary ions that distinguish lignin and polysaccharides in time-of-flight secondary ion mass spectrometry analysis of wood. Anal. Chem. 83, 804–812 (2011).

17. Nakamoto, K. Infrared and Raman spectra of inorganic and coordination compounds. Handb. Vib. Spectrosc. 21 (2006).

18. Huang, W. E., Li, M., Jarvis, R. M., Goodacre, R. & Banwart, S. A. Shining Light on the Microbial World: The Application of Raman Microspectroscopy. Adv. Appl. Microbiol. 70, 153–186 (2010).

19. El-Naggar, M. Y. et al. Electrical transport along bacterial nanowires from Shewanella oneidensis MR-1. Proc. Natl. Acad. Sci. 107, 18127–18131 (2010).

20. Wang, F. et al. Structure of microbial nanowires reveals stacked hemes that transport electrons over micrometers. Cell 177, 361–369.e10 (2019).

21. Franquet, A. et al. Self focusing SIMS: Probing thin film composition in very confined volumes. Appl. Surf. Sci. 365, 143–152 (2016).

22. Horch, M. et al. Resonance Raman spectroscopy on [NiFe] hydrogenase provides structural insights into catalytic intermediates and reactions. J. Am. Chem. Soc. 136, 9870–9873 (2014).

23. Lovering, A. L., Safadi, S. S. & Strynadka, N. C. J. Structural perspective of peptidoglycan biosynthesis and assembly. Annu. Rev. Biochem. 81, 451–478 (2012).

24. Fumagalli, L., Esteban-Ferrer, D., Cuervo, A., Carrascosa, J. L. & Gomila, G. Label-free identification of single dielectric nanoparticles and viruses with ultraweak polarization forces. Nat. Mater. 11, 808–816 (2012).

25. Cuervo, A. et al. Direct measurement of the dielectric polarization properties of DNA. Proc. Natl. Acad. Sci. 111, E3624–E3630 (2014).

26. Fumagalli, L., Ferrari, G., Sampietro, M. & Gomila, G. Quantitative nanoscale dielectric microscopy of single-layer supported biomembranes. Nano Lett. 9, 1604–1608 (2009).

27. Lozano, H. et al. Dielectric constant of flagellin proteins measured by scanning dielectric microscopy. Nanoscale 10, 19188–19194 (2018).

28. Liu, J. et al. Metalloproteins containing cytochrome, iron–sulfur, or copper redox centers. Chem. Rev. 114, 4366–4469 (2014).

29. Boer, J. L., Mulrooney, S. B. & Hausinger, R. P. Nickel-dependent metalloenzymes. Arch. Biochem. Biophys. 544, 142–152 (2014).

30. Can, M., Armstrong, F. A. & Ragsdale, S. W. Structure, function, and mechanism of the nickel metalloenzymes, CO dehydrogenase, and acetyl-CoA synthase. Chem. Rev. 114, 4149–4174 (2014).

31. Desguin, B. et al. A tethered niacin-derived pincer complex with a nickel-carbon bond in lactate racemase. Science 349, 66–69 (2015).

32. Fiedler, A. T., Bryngelson, P. A., Maroney, M. J. & Brunold, T. C. Spectroscopic and computational studies of Ni superoxide dismutase:? Electronic structure contributions to enzymatic function. J. Am. Chem. Soc. 127, 5449–5462 (2005).

33. Bishop, D. W., Thomas, P. S. & Ray, A. S. Micro Raman characterization of nickel sulfide inclusions in toughened glass. Mater. Res. Bull. 35, 1123–1128 (2000).

34. Wang, J.-H., Cheng, Z., Brédas, J.-L. & Liu, M. Electronic and vibrational properties of nickel sulfides from first principles. J. Chem. Phys. 127, 214705 (2007).

35. Faber, M. S., Lukowski, M. A., Ding, Q., Kaiser, N. S. & Jin, S. Earth-abundant metal pyrites (FeS_2_, CoS_2_, NiS_2_, and their alloys) for highly efficient hydrogen evolution and polysulfide reduction electrocatalysis. J. Phys. Chem. C Nanomater. Interfaces 118, 21347–21356 (2014).

36. Johnson, M. K. Vibrational spectra of dithiolene complexes. in Progress in Inorganic Chemistry 52: Dithiolene Chemistry: Synthesis, Properties, and Applications (ed. Stiefel, E. I.) 213–266 (John Wiley & Sons, Inc., 2004). doi:10.1002/0471471933.ch4.

37. Petrenko, T., Ray, K., Wieghardt, K. E. & Neese, F. Vibrational markers for the open-shell character of transition metal bis-dithiolenes: An infrared, resonance Raman, and quantum chemical study. J. Am. Chem. Soc. 128, 4422–4436 (2006).

38. Bostick, C. D. et al. Protein bioelectronics: A review of what we do and do not know. Rep. Prog. Phys. 81, 026601 (2018).

39. Zhang, B. et al. Role of contacts in long-range protein conductance. Proc. Natl. Acad. Sci. 116, 5886–5891 (2019).

40. Malvankar, N. S. et al. Tunable metallic-like conductivity in microbial nanowire networks. Nat. Nanotechnol. 6, 573–579 (2011).

41. Ing, N. L., Spencer, R. K., Luong, S. H., Nguyen, H. D. & Hochbaum, A. I. Electronic conductivity in biomimetic α-helical peptide nanofibers and gels. ACS Nano 12, 2652–2661 (2018).

