## Supplementary info for "Efficient long-range conduction in cable bacteria through nickel protein wires"

**This DOCX file includes:**

Methods
Supplementary Text
Supplementary Figures 1 to 16
Supplementary Tables 1 to 3
Captions for Movies S1

**Other Supplementary information for this manuscript include the following:**

Movie S1
Excel file with Supplementary Tables 2 and 3

### Methods

#### Cultivation and sample preparation

Cable bacteria were cultured in natural sediments which were sieved, homogenized, repacked into PVC core liner tubes (diameter 40 mm) and incubated in aerated artificial seawater at *in situ* salinity (see[1]). Enrichment cultures with sediment collected from a salt marsh creek bed (Rattekaai, The Netherlands) consistently developed thick (~4 μm diameter) filaments that were used for majority of the experiments (unless stated otherwise). To verify whether Raman spectra were similar for different groups of cable bacteria, filaments were retrieved from enrichment cultures using sediments from other environments: Mokbaai (The Netherlands), marine intertidal sediment providing thick filaments (~4 μm), Aarhus university pond (Denmark), a freshwater site with thin filaments (~1 μm); Yarra River, Australia, brackish estuarine sediment, generating thin filaments (~1 μm) that were used for stable isotope labelling.

Under a stereo microscope, cable bacterium filaments were gently pulled from the top layer of the sediment with glass hooks that were custom-made from Pasteur pipettes. Subsequently, cable bacterium filaments were cleaned and washed by transferring them at least six times between droplets (~ 20 μl) of MilliQ water on a microscope cover slip, thus providing "intact cable bacteria filaments". Fiber sheath samples were obtained by extraction of intact filaments in a droplet (~ 20 μl) of 1 % (w/w) sodium dodecyl sulfate (SDS) for 10 min. followed by six MilliQ washes. Subsequently, filaments were incubated for 10 min. in a droplet (~ 20 μl) of 1 mM sodium ethylenediaminetetraacetate (EDTA), pH 8, and again six times washed in MilliQ[2].

#### Confocal Raman microscopy

Raman spectra were recorded on several confocal microscopy systems equipped with different lasers: Renishaw InVia (532 nm green laser, 30 mW), Horiba LabRAM (532 nm green laser,  500 mW) and WITec confocal CRM alpha 300 Raman microscope (30 mW 532 nm green laser and 180 mW 785 nm near infra-red (NIR) laser). Intact cable bacterium filaments and fiber sheaths were deposited on various substrates (glass microscope slides, cover slips, Al-coated Petri dishes) for the green-laser Raman analysis and Raman grade $CaF_2$ covers for combined green-laser and NIR-laser Raman analysis. In addition, we also recorded green laser Raman spectra on living cable bacteria in sediment gradient slides[3]. Raman spectra were subjected to cosmic spike removal (CRR), averaged, and baseline corrected (NIR laser spectra only, using the Asymmetric Least Squares method from the Baseline package in R[4]). The background spectrum - as recorded next to the filaments - was then subtracted.

**Atomic force microscopy – infrared spectroscopy (AFM-IR)**

Fiber sheaths were deposited on gold coated silicon wafers, and AFM-IR spectra were recorded on a NanoIR2 instrument (Anasys). An OPO laser covering the IR range from 900 to 2235 and 2500 to 3600 $cm^{-1}$ (1.3 to 4.7% laser power) was used to record IR spectra. A QCL laser (1500-1700 at 10% laser power) was used to map the Amide I band maximum at 1642 $cm^{-1}$. We recorded AFM images of samples before and after recording the IR spectra to ensure that we did not damage the fiber sheath during the collection of spectra. Contact mode AFM was used throughout with a spring constant $k \sim 0.07$- 0.4 N $m^{-1}$. Tips were supplied by Anasys and are custom made for the NanoIR2. Spectra from both the central cell area and the cell junctions were recorded. Spectra were smoothened with a Savitzky–Golay filter, averaged and the background as recorded next to the filaments was subtracted.

**Scanning Dielectric Microscopy (SDM)**

Scanning Dielectric Microscopy was performed following the methods developed in [5]. Briefly, an AC voltage of amplitude 4 V and frequency 2 kHz was applied between a conductive AFM probe (PtSi-CONT from Nanosensors Gmbh with equivalent spring constant k~0.2 N $m^{-1}$ and resonance frequency $f_0$~13 kHz) of a Cypher S AFM (Oxford Instruments, UK). Fiber sheaths were deposited on highly oriented pyrolytic graphite (HOPG) substrate (μMash Inc.). The amplitude and phase of the 2ω harmonic of the cantilever oscillation were recorded in constant height mode at 0% RH ($N_2$ flow) in SNAP mode. The oscillation amplitude was converted to a capacitance gradient as detailed in[5]. The experimental noise was ~0.7 zF $nm^{-1}$ for the amplitude and ~0.4° for the phase (for more details see[5–7]).

For quantitative analysis through finite-element modelling, we described a periplasmic fiber using the elliptic cylinder model as developed in[6] to which we added a core-shell structure with a conductive core and a surrounding shell of thickness *d*. The fiber dimensions (major and minor semi-axis) were extracted from the de-convoluted topographic images. The accuracy of the elliptic cylinder model was assessed against a more realistic model that includes the small (±5 nm) surface topographic variations of the fiber. This analysis supports the assumption that electrical properties are homogeneous along the longitudinal axis of the fibers (see supplementary text). The core and shell were assumed to have conductivities $\sigma_c$ and $\sigma_s$, and relative permittivities $\varepsilon_c$ and $\varepsilon_s$, respectively. The AFM tip was modelled as a truncated cone ending with a spherical apex[5]. The radius and half cone angle were calibrated from approach curves measured on a bare part of the HOPG substrate[5]. The remaining tip parameters were left to their nominal values (cone height 12.5 μm, disc cantilever thickness 3 μm, cantilever disc radius 3 μm). The macroscopic cantilever contribution was modelled by a constant offset[6]. In the model, a sinusoidal voltage of frequency ω and amplitude $v_{ac}$ is applied between the tip and the metallic substrate. The force acting on the tip (modulus and phase) was calculated by solving the built-in electric currents model, which employs the AC/DC electrostatic module of COMSOL Multiphysics 5.3 with software routines written in Matlab (Mathworks Inc.), as previously described[5–7] (see also Supplementary information).

**Scanning transmission electron microscopy - energy-dispersive X-ray spectroscopy (STEM-EDX)**

Intact cable bacterium filaments or fiber sheaths were deposited on a Formvar-coated Au grid or Quantifoil Cu grid and excess water was removed by filter paper. For STEM-EDX, the grid with the filaments was first dried, whereas for the 3D tomography, the grid was immediately plunged in liquid ethane using a Vitrobot Mark 2 plunge freezer (Thermo Fischer Scientific). Afterwards, grids were transferred to liquid nitrogen (~77 K) and mounted into a Gatan cryo holder. Finally, the sample was freeze dried over a period of 8 h in vacuum at 77 K and subsequently allowed to heat to room temperature[8]. With this sample preparation method, the collapse of the fiber sheath, which happens during conventional drying on a TEM grid, could be prevented. Samples were characterized by high angle annular dark field scanning transmission electron microscopy (HAADF-STEM), electron tomography and energy dispersive X-ray spectroscopy (EDX) using a Tecnai Osiris microscope (Thermo Fischer Scientific) operated at 200 kV. For electron tomography, projection images were acquired over a tilt range of ± 50° with an increment of 2° and reconstructed using a simultaneous iterative reconstruction technique (SIRT) implemented in the Astra toolbox[9]. The EDX measurements were carried out using a ChemiSTEM system and analyzed using the Bruker ESPRIT software.

**Synchrotron low-energy X-ray microscopy and X-ray Fluorescence (XRM and LEXRF)**

Intact cable bacteria and extracted fiber sheaths were deposited on Formvar-coated Au TEM grids and dried. X-ray microscopy and fluorescence experiments were carried out at the TwinMic Beamline[10] in the Elettra Sincrotrone (Trieste, Italy). For the experiments, TwinMic was operated in scanning mode under a microprobe delivered by an Au zone plate diffractive optic with 600 μm diameter and an outermost zone of 50 nm. In this modality, a fast readout CCD camera produces absorption and phase contrast images, thus providing morphological information on the scanned areas. Simultaneously, LEXRF spectra can be acquired by eight silicon drift detectors places in front of the sample, providing the elemental distribution of excited elements. The excitation energy was chosen accordingly to the elements of interest. Here, an excitation energy of 2 keV was used to have optimal excitation of Si, Al, Mg, Fe, Ni and Cu. The beam spot size was 570 nm. Firstly, samples were examined with a light microscope to select the regions of interest, then placed inside the TwinMic vacuum chamber operating at $10^{-6}$ mbar pressure. The acquisition time was typically 20 seconds per pixel. LEXRF maps were obtained by processing the LEXRF spectra with the PyMCA multiplatform software[11].

**Secondary ion mass spectroscopy (SIMS)**

The elemental composition of the fiber sheaths was assessed by nano-scale secondary ion mass spectrometry (NanoSIMS 50L, Cameca) operated at Utrecht University. Upon extraction, fiber sheaths were transferred onto a polycarbonate filter (pore size 0.2 μm, Isopore, Millipore, Netherlands) pre-coated with a 5-nm thick gold layer and air-dried in a desiccator for at least 24h. Fields of view selected with scanning electron microscopy (SEM) were first implanted with low-energy $Cs^+$ ions (<500 eV) to increase the secondary ion yield without sputtering the material of the sample. Subsequently, the high-energy primary $Cs^+$-ion beam (16 keV) was rastered over the sample surface while detecting secondary ions $^{12}C^-$,

$^{16}O^-$, $^{12}C^{14}N^-$, $^{31}P^-$, and $^{32}S^-$. To ensure high lateral (~50 nm) and depth resolution, the primary ion current of 0.5 pA and dwelling time of 1 ms/pixel were used, and the same area (10x10 microns, resolution 256x256 pixels) was imaged multiple times (600 planes total). The magnetic field strength and exact positions of the electron multiplier detectors were adjusted using an element standard (SPI Supplies, 02757-AB 59 Metals & Minerals Standard). The obtained image data was processed with the Look@NanoSIMS software[12].

A similar approach was used for the detection of positive secondary ions $^{56}Fe^+$, $^{58}Ni^+$, $^{60}Ni^+$, $^{63}Cu^+$ and $^{66}Zn^+$ by NanoSIMS, with the following exceptions. Sputtering was done with the primary $O^-$-ion beam (2 pA, beam size ~100 nm) generated by the radio-frequency (RF) plasma source. No low-energy implantation was done with the $O^-$-ion beam. Due to low signals, dwell time was increased to 5 ms per pixel to ensure reliable alignment of the imaged planes; a total of 168 frames were collected. When detecting zinc, isotope $^{66}Zn$ was selected because the position of the detector measuring $^{63}Cu$ was too close to the position required for the measurement of the more abundant isotope $^{64}Zn$.

Time of Flight (ToF)-SIMS (TOF.SIMS 5, IONTOF, Germany) analysis was performed on intact cable bacterium filaments and fiber sheaths to obtain high mass resolution spectra and detailed depth profiles of both organic fragments and metals. A bundle of well washed filaments was deposited on gold-coated silicon wafers. Using incident light microscopy, spots with free lying filaments were identified for ToF-SIMS analysis. The ToF-SIMS was operated in mass spectrometry mode with a $Bi_3^+$ analytical beam (energy 30 keV, current ~0.35 pA, 100x100 $\mu m^2$ area, 256x256 pixels) and an $Ar_{4000}^+$ gas cluster ion beam (energy 10 keV, current ~1 pA, 400x400 $\mu m^2$ area) was interlaced to obtain depth profiles. The SurfaceLab software (IONTOF, Germany) was used for data analysis. Mass spectra were internally calibrated with $C^+$, $CH_3^+$, $C_2H_3^+$, $C_4H_5^+$ and $Au^+$ for positive mode and $CH^-$, $C_2H^-$, $C_3H^-$, $C_4H^-$ and $Au^-$ for negative mode spectra and peaks were identified based on exact mass and isotopic composition for metal ions in combination with the Hybrid-SIMS analysis (see below). Lateral regions-of-interest (ROIs) were selected for the filaments and potential spots of sediment clay particles or minerals (high signals of $S^-$, $P^-$, $Fe^+$, $Ca^+$, $P^+$, $Al^+$ and/or $Si^+$) or salt (high signals of $Na^+$, $K^+$, $Cl^-$) were excluded.

Sputtering depths were calibrated with an *in-situ* AFM that was operated in contact mode. To constrain analysis time, a subarea of either 50x50 $\mu m^2$ or 20x20 $\mu m^2$ (both 256x256 pixels, 8 μm/sec) of the ToF-SIMS analytical area was imaged by AFM. AFM images before analysis and just after the Ni peak or the carbohydrate peak were recorded and aligned and subtracted in the Gwyddion software to obtain average sputtering depths for the fiber sheaths (see Supplementary Figure 10 for an example of the data treatment).

Additionally, we analyzed fiber sheaths with a Hybrid-SIMS (IONTOF, Germany) operated with an $Ar_{4000}^+$ cluster analytical beam and equipped with an Orbitrap mass spectrometer. The advantage of the Orbitrap analyser is the higher mass resolution (~240000) compared to standard ToF-SIMS (~6000 for our samples), and therefore, it was used to identify fragment ions in both positive and in negative modes and to check peak purity. Disadvantages of the Orbitrap are that masses below ca. 50 amu cannot be assessed, which is a minor issue as ToF identification is generally sufficient here, and the detection limit is poorer because of a ca. 200 count background.

To assist with the interpretation of the ToF-SIMS spectra and the identification of the polysaccharide layer, we analyzed additional samples as reference: bovine serum albumin (BSA), starch from potato, pectin from apple, peptidoglycan from *Methanobacterium* (all from Sigma-Aldrich) and a 1:5 mixture of freshly precipitated NiS mineral particles in BSA. These samples were all spotted on gold-coated silicon wafers and analyzed by ToF-SIMS as described above in both positive and negative mode.

**Stable isotope labelling experiments**

Stable isotope labelling was performed by adding $^{13}C$ labelled glucose and $^{34}S$ labelled sulfate to sediments in which cable bacteria were grown. For the $^{13}C$-glucose experiment, 5 mg of glucose was added to each core (42 mm internal diameter) and mixed into the top ~3 cm of the core immediately after it was repacked. These cores were incubated submerged in an aerated aquarium. For the $^{34}S$-sulfate labelling experiment, artificial seawater (ASW) was prepared to a salinity of 20 (to match in-situ salinity) and 20 mM 90 atom % $^{34}S$-sulfate. Before repacking, 25 mL sediment was centrifuged at 110 g for 5 mins (supernatant discarded), re-suspended in 25 mL $^{34}S$-labelled ASW then repacked into 1 cm diameter cut off syringes. Once the sediment had consolidated, the syringes were topped up with $^{34}S$ labelled seawater (~5 cm water column) and a bubbler line was added to each core. Cores were regularly topped up with ultra-pure water to compensate for evaporation. For both labelling experiments a control with unlabeled substrates was treated identically. For $^{13}C$, filaments were collected for Raman imaging at 14 and 21 days after labelling. In the case of $^{34}S$, filaments were collected after 20 and 50 days for Raman imaging.

**The role of the Ni/S group in electron conduction**

Two types of experiments were performed to determine the role of the Ni/S-group in the conduction of the fiber sheath. First, the effect of oxidation or reduction on the Raman signal of the two lower bands at 367 and 492 $cm^{-1}$ was determined. For the Raman measurements, extracted fiber sheaths were pre-treated for 10 min in 20 µl droplets of the reductant (10 mM $K_4Fe^{II}(CN)_6$) or oxidant (10 mM $K_3Fe^{III}(CN)_6$) and enclosed in the same solute between a glass microscope slide and coverslip sealed with nail polish. At least 15 spectra from 5 individual fiber sheaths were recorded with a 532 nm green-laser Raman microscope as described above. Subsequently, the effect of oxidation or reduction on the conductance of fiber sheaths was evaluated. To this end, individual fiber sheaths were stretched out on a microscope cover slip and a small spot of conductive water-based carbon paste (EM-Tec) was added as an electrode on both ends of the filament. After the carbon paste electrodes had dried, they were connected with conductive copper tape and additional carbon paste to a Palmsens 4 potentiostat and sealed with nail polish to make them water resistant. The effect of reduction or oxidation on filament conductance was determined by adding 2 µl of 10 mM $K_4Fe^{II}(CN)_6$) or 10 mM $K_3Fe^{III}(CN)_6$ respectively to the fiber sheath area between the electrodes. As control we added a 2 µl droplet of MilliQ. The electrical current was measured directly before and after drop addition. Adding a second (i.e. fresh) 2 µl droplet of oxidant or reductant typically did not result in a further change in conduction. We performed repeated oxidation, reduction and MilliQ cycles on single filaments. Occasionally unexpected, sharp drops in conduction were seen between treatments, suggesting the fiber sheath had been damaged during exchange of solutes. These suspect data were excluded from the analysis. Conductance measurements were performed at room temperature and current - voltage (IV) curves were recorded (bias -0.1 to 0.1 V, scan rate 0.01 $Vs^{-1}$). The length ($\Delta x$) of the filament between the electrodes was measured with a Dino-lite microscope. Normalized current was calculated as $I_{norm} = I*(\Delta x/ \Delta x_{ref})$ with $\Delta V = 0.1$ V and $\Delta x_{ref} = 0.3$ mm (see[13]).

In a second set of experiments, Ni was removed selectively by extraction in 50 mM EDTA and effects on Raman signals and conduction were determined. For the Raman measurements, SDS-only extracted fiber sheaths were treated for 10 or 40 min in 20 µl droplets of 1 or 50 mM EDTA and deposited after 5 MilliQ washes on a glass microscope slide. At least 15 spectra from 5 individual fiber sheaths were recorded with a 532 nm green laser as described above. Additionally, the conductance of 10 individual extracted fiber sheaths that were either treated with the standard 10 min 1 mM EDTA or 10 min 50 mM EDTA were compared. Conductance was measured in terms of normalized current as described above. A 10 min extraction time was used for both treatments as this resulted in similar handling times. Fresh EDTA solutions were prepared daily from 0.5 M EDTA stock solution (pH 8). Morphology of the fiber sheaths treated with 1 mM or 50 mM EDTA was studied by AFM as described in[2].

### Supplementary text

#### Additional results and discussion of the ToF-SIMS analysis

The principal results of the ToF-SIMS analysis are described in the main text. Here we provide additional experimental results and discussion to support the conclusions drawn in the main text.

*Fragment identification*. In total 71 fragment ions were identified in positive mode and 173 in negative mode in the ToF-SIMS runs of the fiber sheaths. Identification was based on the exact mass in ToF-SIMS analysis and was checked for identity and purity with the Orbitrap-SIMS runs. For transition metals and S, the isotopic mass distribution was verified. For Ni, the four main isotopes ($^{58}$Ni, $^{60}$Ni, $^{61}$Ni and $^{62}$Ni) were found with the expected isotopic abundance, whereas the minor isotope $^{64}$Ni showed mass interference from a low amount of Zn. For Cu, both isotopes ($^{53}$Cu and $^{65}$Cu) showed the expected abundance. For Fe, the two main Fe isotopes ($^{54}$Fe and $^{56}$Fe) showed the expected abundance, whereas the minor Fe isotopes ($^{57}$Fe and $^{58}$Fe) showed mass interference from an unknown source and Ni respectively. Zn counts were was low and not reliable as the main isotope ($^{64}$Zn) showed mass interference from $^{64}$Ni. Other transition metals found in metalloproteins like Mn, Co, Mo and W were not detected or had very low counts. The main isotopes of Sulfur ($^{32}$S and $^{34}$S) also showed the expected abundances, whereas the minor isotopes $^{33}$S and $^{36}$S showed mass interference from an unknown source. Assignment of the identified fragment ions to biochemical components of intact cable bacteria and the fiber sheath was based on literature[14–20] and specific depth profiles (see Supplementary Table 2 and 3, Fig. 1, and Supplementary Figure 1).

*Transition metals.* The ToF-SIMS depth profiles of Ni and Fe are discussed in detail in the main text. Cu showed a peak at the first data point of the depth profile in both the fiber sheaths and the intact cable bacteria (Fig. 1D, Supplementary Figure 1A-B, Supplementary Figure 11), though with a highly variable count number. This surface peak and its variability between samples was consistent with our LEXRF analysis (Fig. 3), which also showed a highly variable Cu signal. However, Cu counts from STEM-EDX analysis and the majority of the ToF-SIMS depth profiles in the fiber sheaths were low compared Ni counts. This suggests that Cu is not present in high concentrations, but likely derives from an impurity that adsorbs to the sample surface at some stage during sample preparation (either cable bacteria collection, fiber sheath extraction procedure) or analysis.

*Polysaccharide fragments.* As discussed in the main text, the basal polysaccharide layer (Fig. 1D and 1E) most likely consists of peptidoglycan, which consists of an acidic amino sugar backbone interconnected by short peptides[21]. We therefore expected to find fragments that contain both oxygen and nitrogen, as derived from the amino sugar backbone of peptidoglycan. To verify that we can detect peptidoglycan specific fragments among these O- and N rich fragments, we performed ToF-SIMS analysis of four protein and polysaccharide reference samples: Bovine Serum Albumin (BSA), starch, pectin, and peptidoglycan (Supplementary Figure 8). Although the mass spectra of these reference samples showed major differences in both negative and positive mode (Supplementary Figure 8), we could not detect any peptidoglycan specific fragments in the fiber sheaths. Moreover, fragments that contained both nitrogen and oxygen showed either the same depth profile as amino acid (protein) derived fragments (i.e. an initial peak at the start of the depth profile), or were specific for nucleic acids that showed low counts and variable depth profiles (Supplementary Table 2 and 3). The latter suggests that some residual DNA/RNA remains associated with the fiber sheaths after extraction. The lack of peptidoglycan specific fragments is most likely due to the abundance of protein in the fiber sheath, which leads to high signals of amino acid derived fragments that mask fragments derived from peptidoglycan. Further work is needed to confirm that the polysaccharide layer is indeed made of peptidoglycan.

*P-containing fragments.* The fiber sheaths also showed high counts of various P-containing fragment ions in both positive and negative mode (Supplementary Table 2 and 3) with variable depth profiles between runs. The likely source of these P-containing ions is the poly-phosphate granules that are commonly found in cable bacteria[22]. Likely, the poly-phosphate granules are incompletely removed during the fiber sheath extraction procedure, and they comprise the particles that are seen in the interior of fiber sheaths during STEM 3D-tomography (Fig. 1A).

*S and Ni containing fragments in negative mode.* Negative mode ToF-SIMS depth profiles showed a subsurface peak of various S-derived anions ($^{32}S^-$, $^{34}S^-$, $SH^-$ and $S_2^-$) at approximately the same position as the Ni peak in positive mode (Fig. 1D and 1E, Supplementary Figure 1, Supplementary Table 3). In addition, a substantial number of other Ni-containing fragments were found in negative mode that also showed this distinct subsurface peak (Fig. 1D, Supplementary Figure 1, Supplementary Table 3). The most prominent Ni containing fragments were a series of $Ni_xS_y^-$ cluster ions ($x = 1$ to 6, $y = x +/- 1$, both $^{58}Ni$ and $^{60}Ni$ isotopologues were detected, Supplementary Figure 9). Additionally, we found $NiC_xN_x^-$ ($x = 1$ or 2) and $NiCSN^-$, and even a small but detectable $Ni^-$ signal (Supplementary Table 3). All these Ni-containing fragments showed a similar sharp subsurface peak in the depth profile as found for Ni in positive mode (Fig. 1E). The observed NiS clusters are most likely formed in the ion plume of the ToF-SIMS rather than being natively present in the fiber sheath[23–25]. As a control, we analyzed an artificial mixture of protein (BSA) and freshly precipitated mineral NiS, and we indeed observed the same $Ni_xS_y^-$ cluster ions (Supplementary Figure 9D). Still, due to an effect called self-focusing[23], the detection of NiS clusters suggests that Ni and S must be present in close proximity in the fiber sheath (lateral (XY) within <0.5 nm, depth (Z) within <2 nm), otherwise they would not be formed in the ToF-SIMS ion plume. This observation is hence in agreement with the presence of a S-ligated Ni cofactor in the fiber sheaths.

We also found a number of organic S fragments in negative mode ToF-SIMS spectra (Fig. 1E, Supplementary Figure 1, Supplementary Table 3). Most of these fragments showed a similar depth profile as $S^-$ with a sharp subsurface peak on top of a general S signal, and so again, we cannot exclude that they were formed during ToF-SIMS analysis. However, two fragments ($C_2S_2^-$ and $C_2S_2H^-$) showed specific depth profiles, with the subsurface peak much more pronounced than for $S^-$ (Fig. 1E). This suggests that these two fragments originate directly from the fiber sheath. These fragments could indicate that the fiber proteins are rich in disulfide bridges (C-S-S-C) or they could be derived from the Ni/S group. Fiber sheaths display an exceptional chemical resistance, as they remain their integrity and conductivity after SDS and EDTA treatments, and this resistance could be aided by protein disulfide bridges.

*Sputtering depth calibration.* Sputtering depth was calibrated by *in-situ* AFM height measurements at three different times (Supplementary Figure 2): before sputtering, just after the Ni or S peal appeared, and finally after the carbohydrate peak appeared. Sputtering depth was approximately linear with sputtering time, although the initial protein layer seemed to sputter somewhat faster (Supplementary Figure 2). A representative example of the resulting AFM height maps is shown in Supplementary Figure 10, where sputtering was stopped at 57 sec (i.e. just after the Ni peak had emerged). There was considerable lateral variation in sputtering depths for a given sputtering time as seen in transects across the central cell area (Supplementary Figure 10D) and at the cell junctions (Supplementary Figure 10C). As the junctions are higher than the cell areas (e.g. Supplementary Figure 10A), the shading in sputtering seen at the cell junctions probably due the angled Ar-cluster sputter beam hitting the side of the cell junction facing the beam at a higher angle whereas the opposite side is hit much less. This lateral variation in sputtering rate together with the left-over cytoplasm content and the presence of the cart-wheel structure at the junctions leads to smearing of the ToF-SIMS depth profiles. This explains why only the first fiber sheath layer is clearly seen in the ToF-SIMS depth profiles.

*Intact cable bacteria.* ToF-SIMS depth profiles from intact cable bacteria also show subsurface peaks for Ni and S containing fragments (Supplementary Figure 11), thus providing additional proof for the Ni and S rich nature of the periplasmic fibers. Comparing the relative position of these fragments to fatty-acid fragments derived from membranes shows that the Ni and S containing fragments originate from the periplasmic space (Supplementary Figure 11), thus confirming that the Ni- and S-rich fibers are embedded in the periplasmic space.

**Additional results and discussion of the SDM analysis**

The key results of the Scanning Dielectric Microscopy (SDM) analysis are described in the main text. Here we describe additional results that strengthen the conclusions obtained. The finite element model we used is the same as presented earlier[5–7] except that here, fibers have a core-shell structure with a conductive core and an insulating outer shell. The whole simulation domain (which encompasses the ellipsoid fiber and the surrounding space) is cylindrical with height 34 μm and radius 17 μm. Insulating boundary conditions are assumed on the lateral and top borders of the simulation domain. A cross-section of the electric potential distribution obtained from the model calculations is shown in the right insert in Fig. 5C. At the frequency of the calculations (2 kHz) and for the parameters in Fig. 5, the electric potential is real, and hence, the phase of the electric potential is constant in space. The length of the fiber in the model is L =1 μm. It has been shown previously that for L >100 nm the force acting on the tip is independent from the fiber length[6].

Supplementary Figure 12A shows experimentally derived capacitance gradient cross-section profiles corresponding to SDM images (similar to the one shown in the left insert in Fig. 5B of the main text) acquired at different tip-substrate distances (continuous lines, ranging from z=60 nm to z=240 nm). The profile at z=66 nm is shown in Fig. 5C of the main text. The dashed lines represent the result of the theoretical calculations with the model described above for the same fiber parameters as those use in Fig. 5 of the main text (h=42 nm, w=87 nm, d=12 nm, $\varepsilon_s = \varepsilon_c = 3$, $\sigma_s = 0$ S/cm, $\sigma_c = 20$ S/cm). The tip-substrate distances have been determined from a capacitance gradient approach curve measured on a bare part of the substrate (Supplementary Figure 12B, symbols) following procedures previously reported[5]. The calculated capacitance gradient profiles nicely reproduce the experimental ones with no adjustable parameter. At all distances, the theoretical (and experimental) electric force phase contrast is zero (not shown). Similar conclusions are reached if we analyze the capacitance gradient approach curves measured on the substrate and on the fiber (pink and black thick lines in Supplementary Figure 12B). A least square fitting of the model curves to the experimental curve for $\varepsilon_s = \varepsilon_c = 3$ and $\sigma_s = 0$ S/cm, $\sigma_c = 20$ S/cm gave d=12±2 nm (red continuous line in Supplementary Figure 12B), in agreement with the value obtained from the capacitance gradient cross-section profile analysis. The sensitivity of the capacitance gradient profiles on the thickness of the insulating shell is shown by the dashed lines in Supplementary Figure 12B for the approach curves and in Supplementary Figure 12C for the profiles at z=66 nm. In these figures we also compare the predictions corresponding to an homogeneous conductive model, corresponding to d=0 nm (or to $\sigma_s = \sigma_c = 20$ S/cm, dark grey line), and to an homogeneous insulating model, corresponding to d=h/2 (or $\sigma_c = 0$ S/m, light grey line). In all cases we assumed a protein composition of all layers, $\varepsilon_s = \varepsilon_c = 3$. The pure conductive model overestimates the force acting on the tip, while the pure insulating model underestimates it.

We have considered other possible sets of electric parameters for the fiber to explore alternative interpretations of the SDM results. Figs. S13A and S13B show, respectively, the contrast values of the amplitude (in zF/nm) and phase of the electrical force for the tip at the center of the fiber at a distance z=66 nm from the substrate as a function of the conductivity of the core in the range from $\sigma_c = 10^{-9}$ S/m (insulator) to $10^3$ S/m (conductor). The dielectric constants have been fixed to $\varepsilon_s = \varepsilon_c = 3$ (proteins). The thickness of the insulating layer has been varied from d=0 nm to d=20 nm. The calculations were done for the frequency of the experiments (2 kHz). The amplitude as a function of the core conductivity displays two plateaus for low and high conductivities, separated by a transition region, which tends to show a third plateau not fully displayed. The phase shows a minimum for every transition region mentioned above. Such behavior is characteristic of materials with (equivalent) permittivities showing real (dielectric) and imaginary (conductive) parts. An analytical expression for the equivalent homogeneous permittivity of the core-shell cylinder in a non-uniform electric field cannot be derived, but the behavior observed is qualitatively similar to the one corresponding to a core-shell spherical particle in a uniform electric field[26]. For a given measured capacitance gradient contrast (grey band in Supplementary Figure 13A), one can obtain couples of values for the shell thicknesses and core conductivity that match[5,27]. For instance, we observe that the solution found above for the shell thickness (d=12 nm) is valid for $\sigma_c > 10^{-2}$ S/m (and, in particular, for $\sigma_c = 2 \cdot 10^3$ S/m as employed here). Other couples of values are for instance d=5 nm and $\sigma_c = 5 \cdot 10^{-6}$ S/cm or d=0 nm, $\sigma_c = 7 \cdot 10^{-7}$ S/cm, among others. For these values the same capacitance gradient profile (matching the experimental one) is obtained, as shown in the Supplementary Figure 13C. However, the different couples of values predict different electrical phase contrast profiles (see Supplementary Figure 13D). Only the couple of values corresponding to the solution reported in the main text (d=12 nm and $\sigma_c > 10^{-2}$ S/m) predicts a null phase contrast, as seen in the experimental results.

We have also considered a homogeneous dielectric model, with no conductivity, and analyzed the (equivalent) homogeneous dielectric constant that the fiber should have to explain the experimental results. A least square fitting of the calculated capacitance gradient approach curves for different dielectric constants to the experimental curve measured on the fiber gives $\varepsilon_s = \varepsilon_c = 11 \pm 3$ (see Supplementary Figure 14A). For this equivalent dielectric constant value the constant height capacitance gradient profiles at the different tip substrate distances adequately reproduce the experimental ones (Supplementary Figure 14B). The phase contrast for this model is zero (not shown). However, a problem is that the equivalent dielectric constant value found ($\varepsilon_s = \varepsilon_c = 11 \pm 3$) is much larger than values obtained, with the same technique, on other (dry) bio-samples made of lipids ($\varepsilon$=2)[28], proteins ($\varepsilon$=3–5)[27,28,6,7] and even nucleic acids ($\varepsilon$~8.5)[5,27] (see Supplementary Figure 14C). Our composition data all suggest that the fibers are made primarily of protein. During ToF-SIMS analysis of fiber sheaths, some fragments derived from residual RNA or DNA were detected, but these were not associated with the initial fiber protein layer (see Additional results and discussion of the ToF-SIMS analysis) and it therefore seems unlikely that an isolated fiber as analyzed here by SDM would contain major amounts of DNA. For this reason, the pure dielectric model is discarded in favor of the conductive core-shell model.

We have also analyzed the homogeneity of the fiber electrical properties along its longitudinal direction in view of the slight variations of the electric contrast observed in the SDM images along the fiber (see left insert in Fig. 5C of the main text). To this end we considered the core-shell geometrical model that accounts for the observed (tiny) height variations of the fiber height (±5 nm)[29] shown in Supplementary Figure 15A. Examples of calculated longitudinal and transversal electric potential distributions are shown in figs. S15B and S15C, respectively. The tip and electrical parameters of the fiber are the same as those used in the calculations of in Fig. 5 of the main text, except for the shell thickness, which here is d=10±2 nm. The slightly smaller value obtained, as compared to d=12±2 nm for the cylindrical model, is due to the tip convolution effects included in this geometrical model (see Supplementary Figure 15D), as discussed elsewhere[29]. Supplementary Figure 15E shows a calculated constant height SDM image at z=66 nm corresponding to the region enclosed by the dashed rectangle in the insert (which corresponds to the image in the left insert in Fig. 5C of the main text). The calculated SDM image is nearly identical to the experimental one, a fact that is further evidenced by comparing the capacitance gradient cross-section profiles calculated on the hills and valleys of the image with the experimental ones (figs. S15F and S15G, respectively). This result shows that, to a good approximation, the electrical properties of the fiber are homogeneous along its length.

Finally, we have also analyzed the electric properties of the fibers when they are still embedded within the fiber sheath by SDM. Supplementary Figure 16A shows an AFM topographic image of a fiber sheath. Figs. S16B and S16C show, respectively, topographic and constant height SDM images acquired in the area enclosed by the dashed rectangle in Supplementary Figure 16A. The corresponding height and capacitance gradient profiles along the dashed lines in figs. S16B and S16C are shown in figs. S16D and S16E, respectively (black lines). The geometry of the tip has been calibrated with a capacitance gradient approach curve acquired on the substrate (not shown) giving R=26±2 nm, θ=22±3º, $C'_{offset}$=109±3 zF/nm. The sample geometry has been reconstructed by using a topographic reconstruction procedure[29], as above (see Supplementary Figure 16D). For simplicity, we have considered an equivalent homogeneous dielectric model characterized by an (equivalent) dielectric constant $\varepsilon_{sheath}$. An example of a calculated electric potential distribution is shown in

Supplementary Figure 16F for a tip-substrate distance z=155 nm and $\varepsilon_{sheath}$=7. Supplementary Figure 16E shows calculated constant height capacitance gradient profiles along the dashed line in Supplementary Figure 16C for different values of $\varepsilon_{sheath}$ (thin lines). The experimental values on the hills (corresponding most likely to fiber positions) agree with the calculated ones for $\varepsilon_{sheath}$~7−11 (except the first hill that gives a somewhat larger value). These values are in reasonable agreement with those found for the isolated fiber when the equivalent dielectric model is considered ε~11±3. This result implies that the electrical properties obtained for the fibers when within the fiber sheath are consistent with those obtained on isolated fibers. Similar results are expected for a fiber sheath model that included its internal structure, composition and electrical properties (e.g. conductive core-shell model for the fibers), as the one sketched in Fig. 5A of the main text. However, obtaining quantitative predictions from SDM measurements for such complex model involving buried structures lies outside the current capabilities of SDM[30] since too many unknowns are present in the model (e.g. fiber position within the fiber sheath, number of fibers present within the polysaccharide layer, thickness of the polysaccharide layer, etc.).

### Supplementary figures and tables

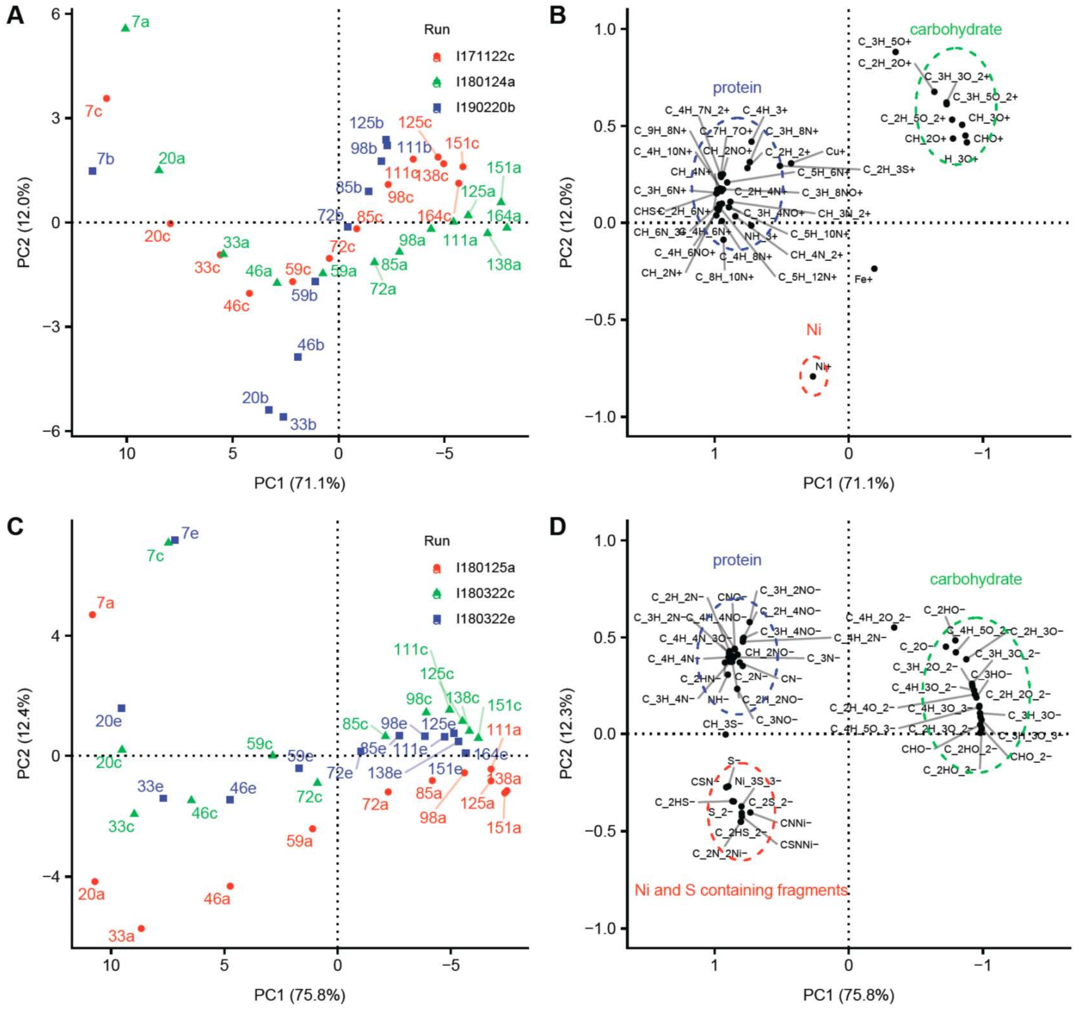


**Supplementary Figure 1.** Principal Component Analysis (PCA) of ToF-SIMS depth profiles for a selection of ion fragments recorded from fiber sheaths of cable bacteria. ToF-SIMS data were collected in both positive (A, B) and negative modes (C, D), and the results of three replicate depth profiling runs are shown. Counts for fragments were normalized and centered. Panels (A, C) show the scores plot (labels indicate sputtering time in seconds; colors denote separate runs). Panels (B, D) show the loading plots of the fragments. Only the data until the peak carbohydrate-derived fragments at 125-164 seconds of sputtering time were retained (representing the first half of the double-folded fiber sheath).

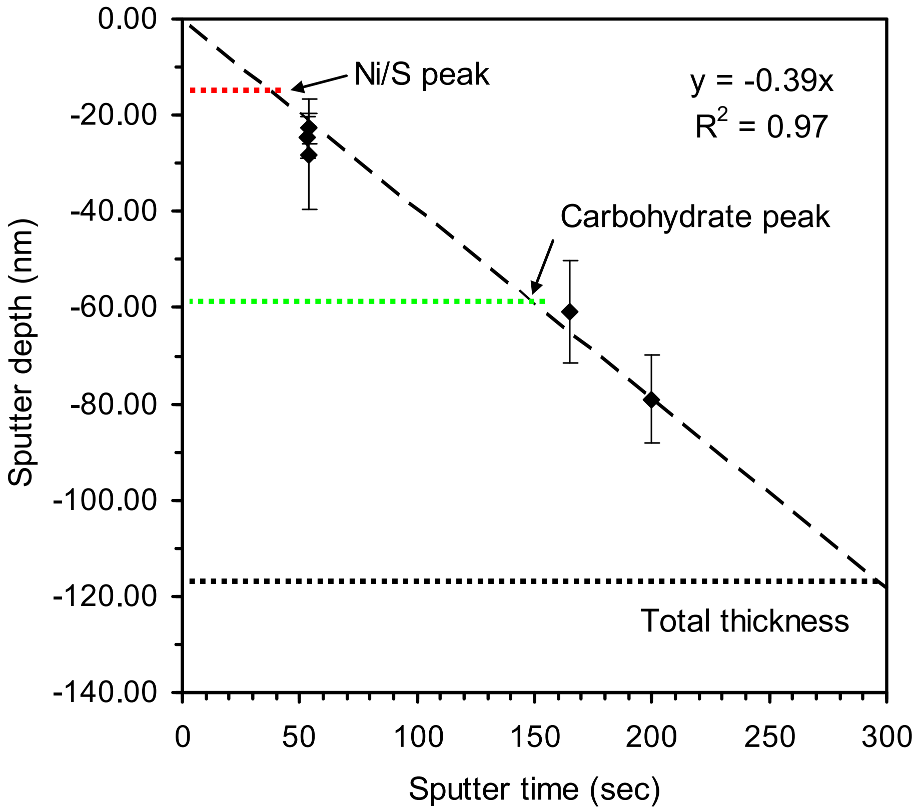


**Supplementary Figure 2.** Calibration of sputtering depth as a function of sputter time during ToF-SIMS analysis. The in-situ AFM within the ToF-SIMS instrument was used to record height images at three different times (start (sputter time = 0 sec), just after the Ni or S peak (ca. 50 sec) and just after the carbohydrate peak (ca. 160 and 190 sec)). Sputtering depth showed a linear relation with sputtering time (dashed line = regression line through origin), though the initial protein layer seemed to sputter somewhat faster. The total thickness of the double-folded fiber sheath amounts to 117 ± 10 nm (determined within the middle of cells at t = 0 sec; black dotted line) and is well in agreement with previous independent AFM imaging [2]. The average position of the Ni/S peak (15 ± 3 nm) and the carbohydrate peak (59 ± 6 nm) are indicated (red and green dotted lines respectively, N = 6 from 3 positive and 3 negative depth profiles in Supplementary Figure 1).

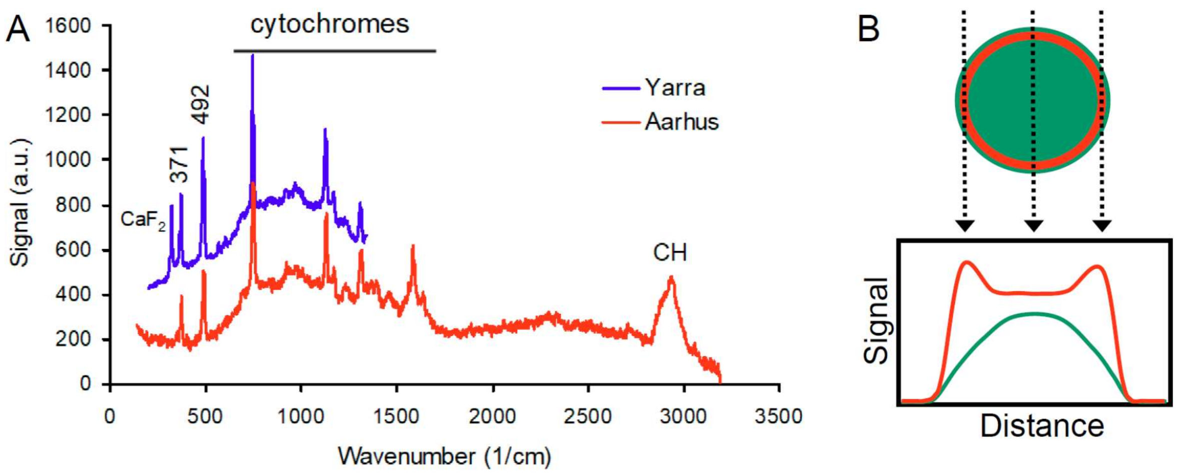


**Supplementary Figure 3.** (A) Green-laser Raman spectra of intact cable bacteria from additional sediments also show the characteristic low-frequency bands at 371 and 492 $cm^{-1}$. Spectra are from ca. 1 µm wide freshwater cable bacteria (Aarhus pond sediment, intact living filaments in glass micro-chambers, average background-corrected spectrum) and ca. 1 µm wide brackish cable bacteria (Yarra River sediment, intact filaments air-dried on a $CaF_2$ cover, average raw spectrum). (B) Interpretation of the Raman signal distribution in the scan across an individual cable bacterium in Fig. 2B. General components of a bacterial cell such as proteins and CH are more-or-less evenly distributed in the cross section (green filled circle), whereas the Ni/S group is only found in the periplasmic space (red open circle). In the depth integrated Raman signals, this leads to a unimodal distribution for Raman bands from the general components and a bimodal distribution for the two low frequency Ni/S bands with a peak at both edges of the filament.

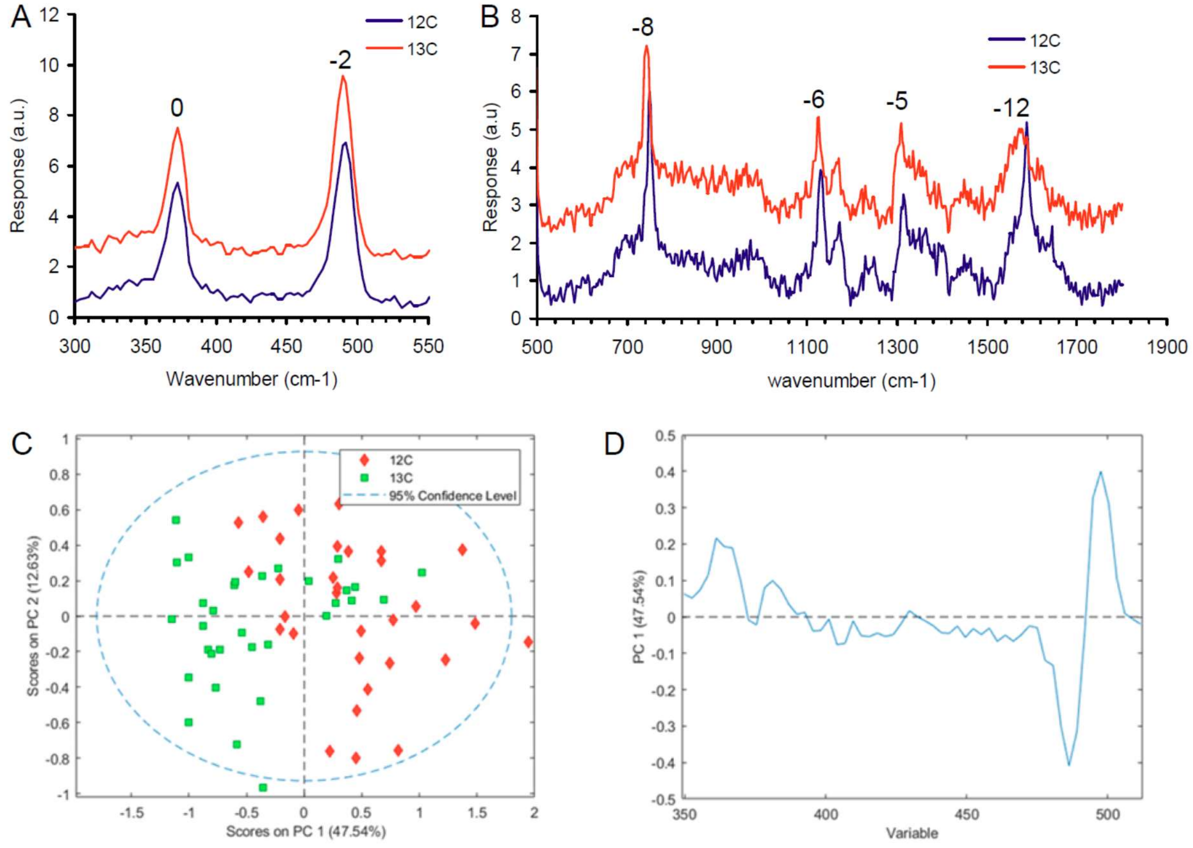


**Supplementary Figure 4.** Effect of $^{13}C$ labelling on the green-laser Raman spectra from intact cable bacteria. A) The low-frequency bands either do not shift (371 cm$^{-1}$ band) or are only slightly affected (491 cm$^{-1}$ band shifts to 489 cm$^{-1}$) by the labelling. B) The characteristic cytochrome bands clearly shift to lower values suggesting that cable bacteria filaments were highly labelled with $^{13}C$. The average of N = 32 spectra is shown for both unlabeled control and $^{13}C$ treatments and shifts in wavenumbers are indicated. C) PCA analysis of the 350 to 550 cm$^{-1}$ region of the spectra containing the low-frequency bands shows a difference between the labelled versus unlabeled treatments. D) Variable (wavenumber) loadings on the first PCA axis show that only the second lower band at 492 cm$^{-1}$ shifted slightly to lower values in the labelled spectra.

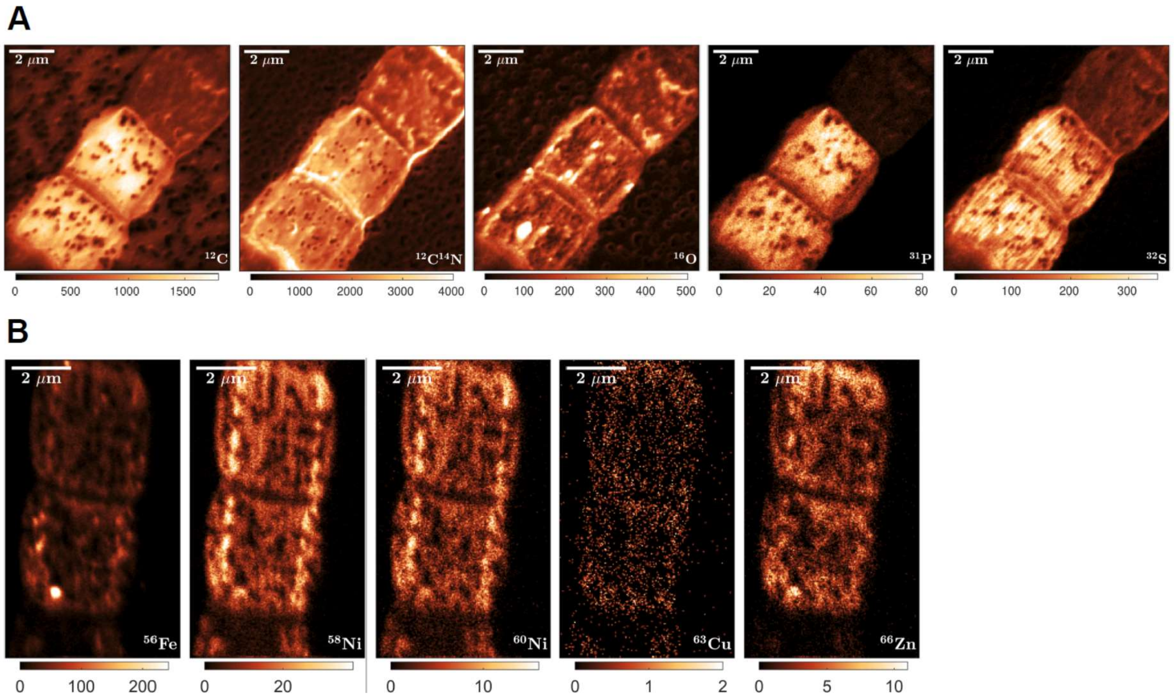


**Supplementary Figure 5.** NanoSIMS images of A) negative ions and B) positive ions of metals from single fiber sheaths deposited on a gold-coated polycarbonate filter. In A) only the first 100 planes of analysis were selected as this most clearly showed the fibers structure in the $^{32}S$ image. In B) the first 50 planes were selected as this showed the fibers most clearly in the $^{58}Ni$ image.

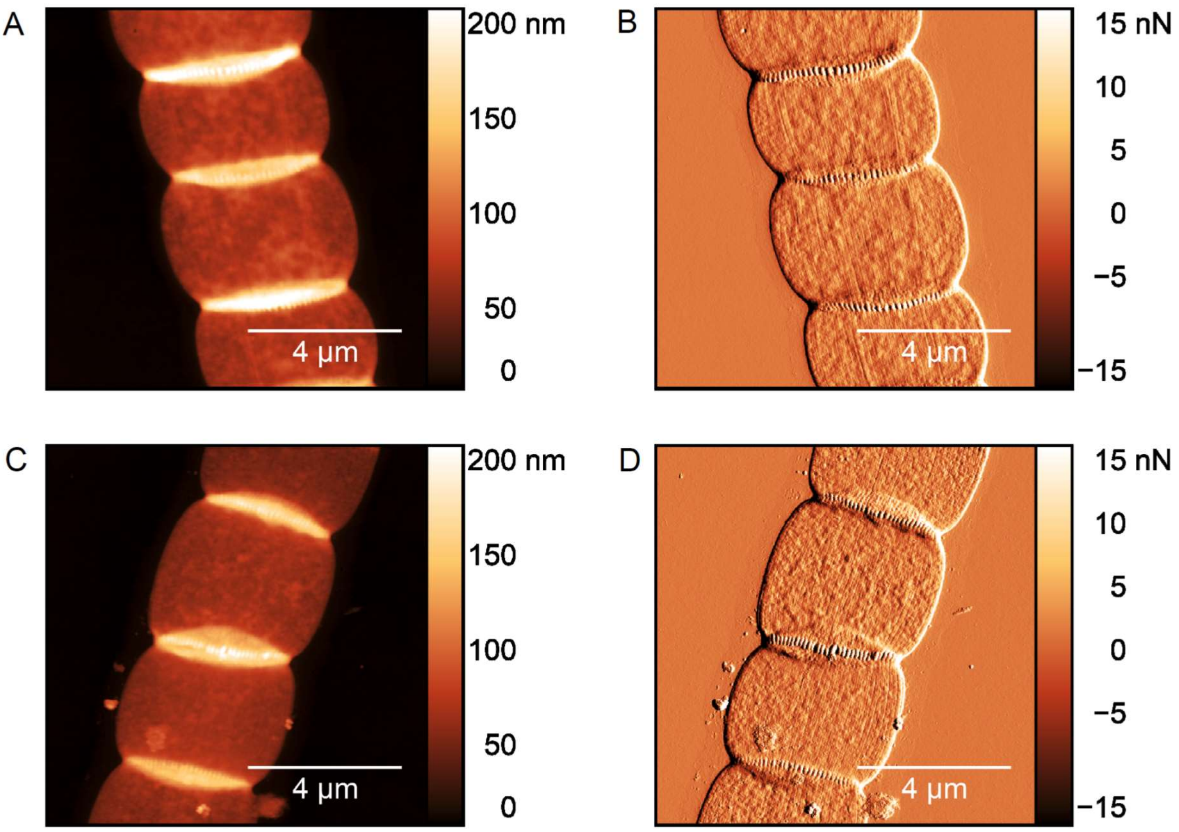


**Supplementary Figure 6.** Atomic Force Microscopy (AFM) images of fiber sheaths as extracted with the standard protocol (A and B, SDS+1 mM EDTA for 10 min) and the high EDTA treatment to remove Ni (C and D, SDS+50 mM EDTA for 10 min). Shown are the height (A,C) and peak force error (B,D) data. The parallel fiber structures are retained after high-concentration EDTA extraction.

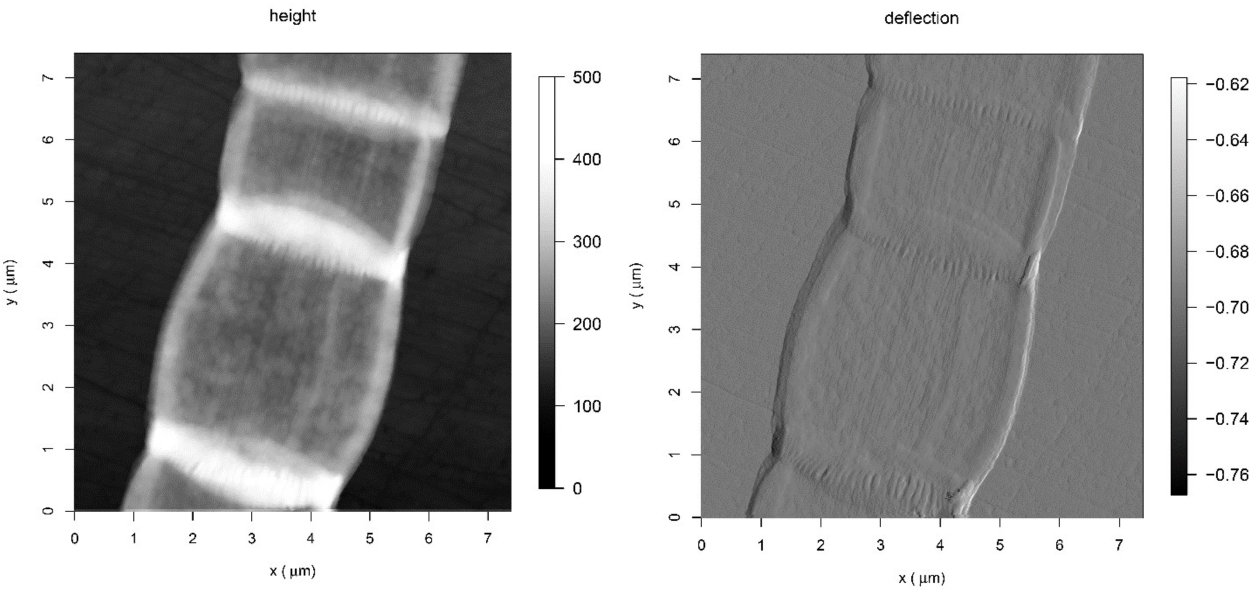


**Supplementary Figure 7.** AFM-IR height (in nm) and deflection maps of extracted fiber sheath showing the parallel fibers.

A

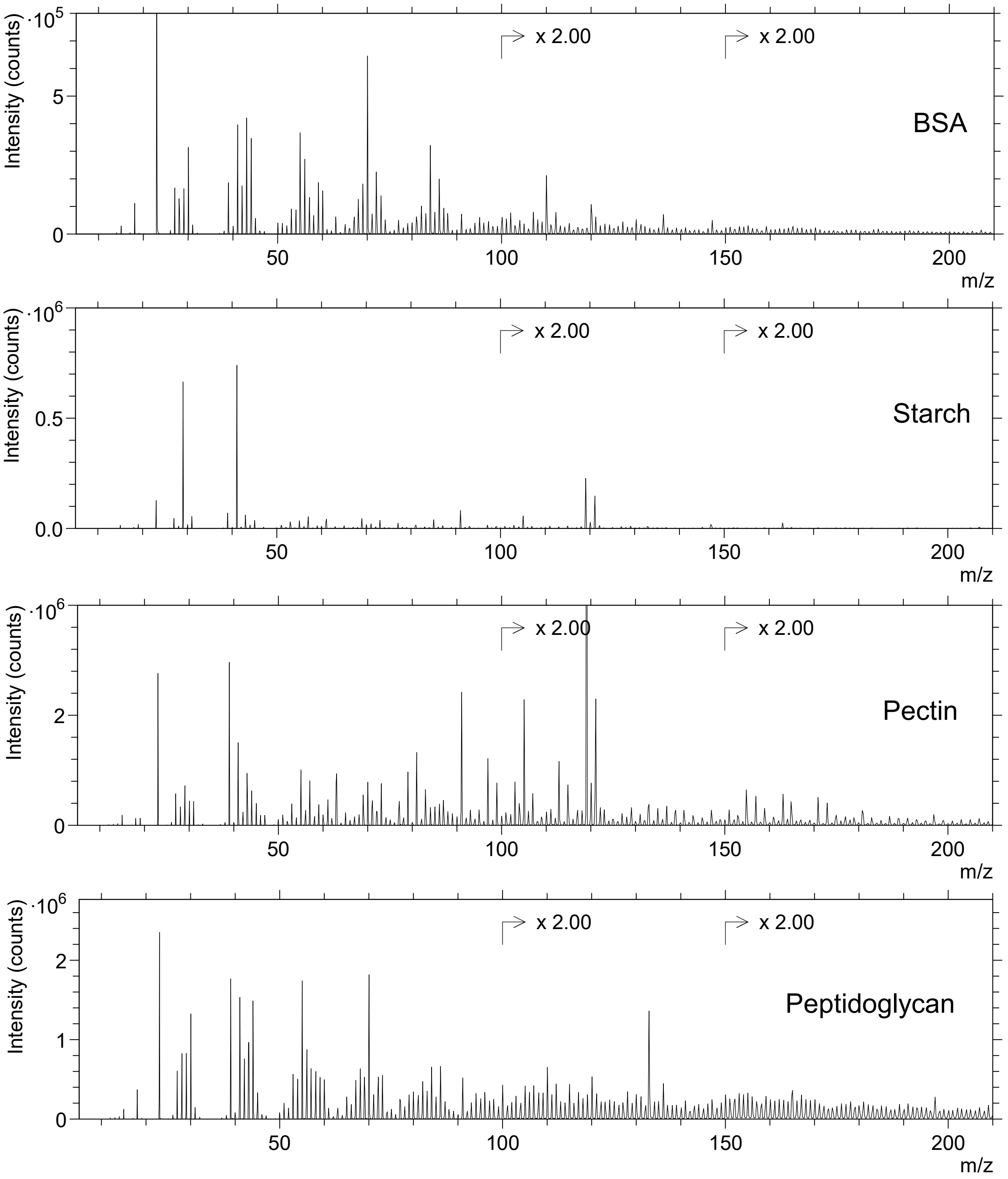
BSA
Starch
Pectin
Peptidoglycan
Intensity (counts)
m/z
x 2.00
·10^5
·10^6
50
100
150
200

563

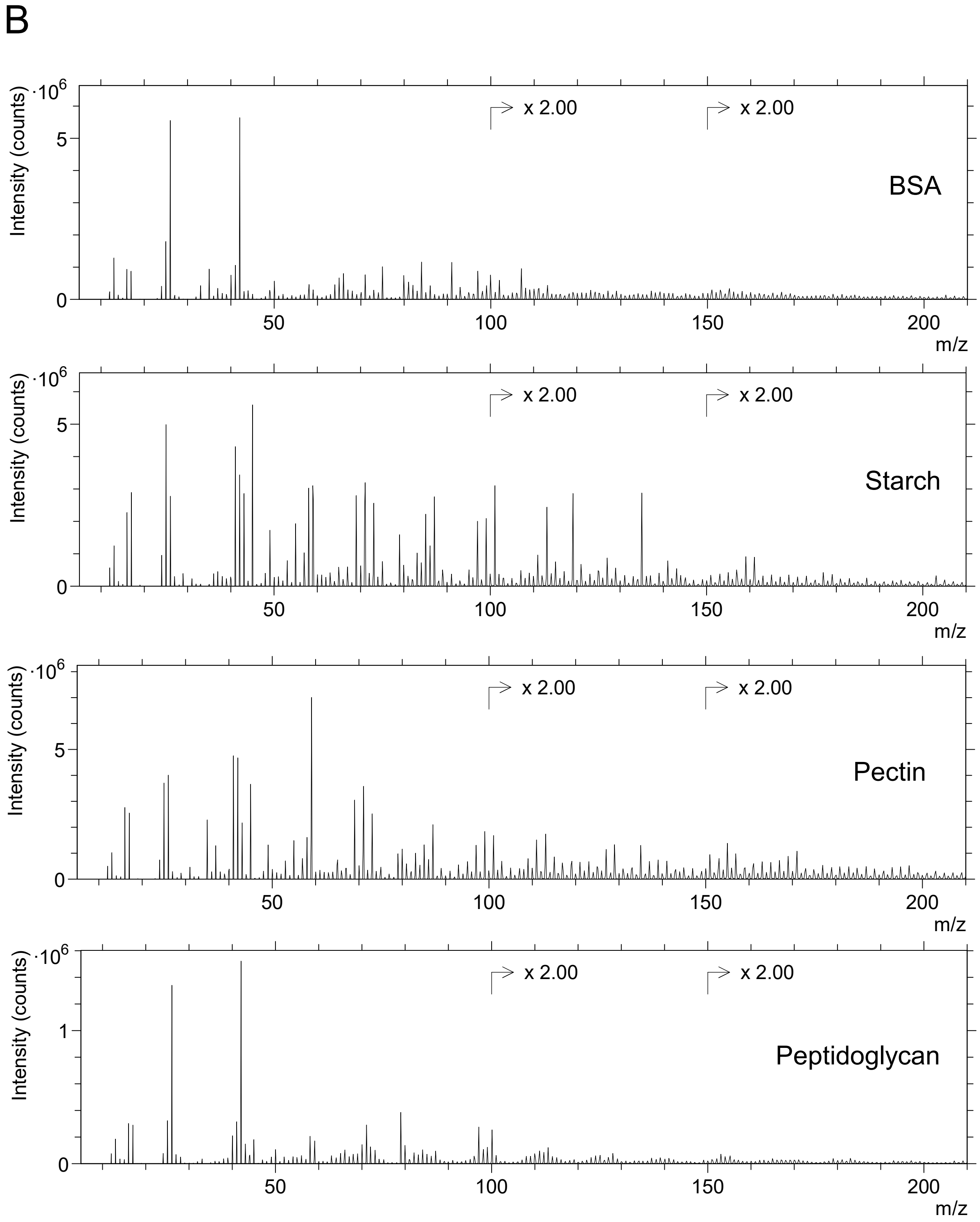


**Supplementary Figure 8.** ToF-SIMS positive (A) and negative (B) mode mass spectrum of additional protein and polysaccharide reference samples analyzed to aid in the interpretation of the fiber sheath mass spectra.

A

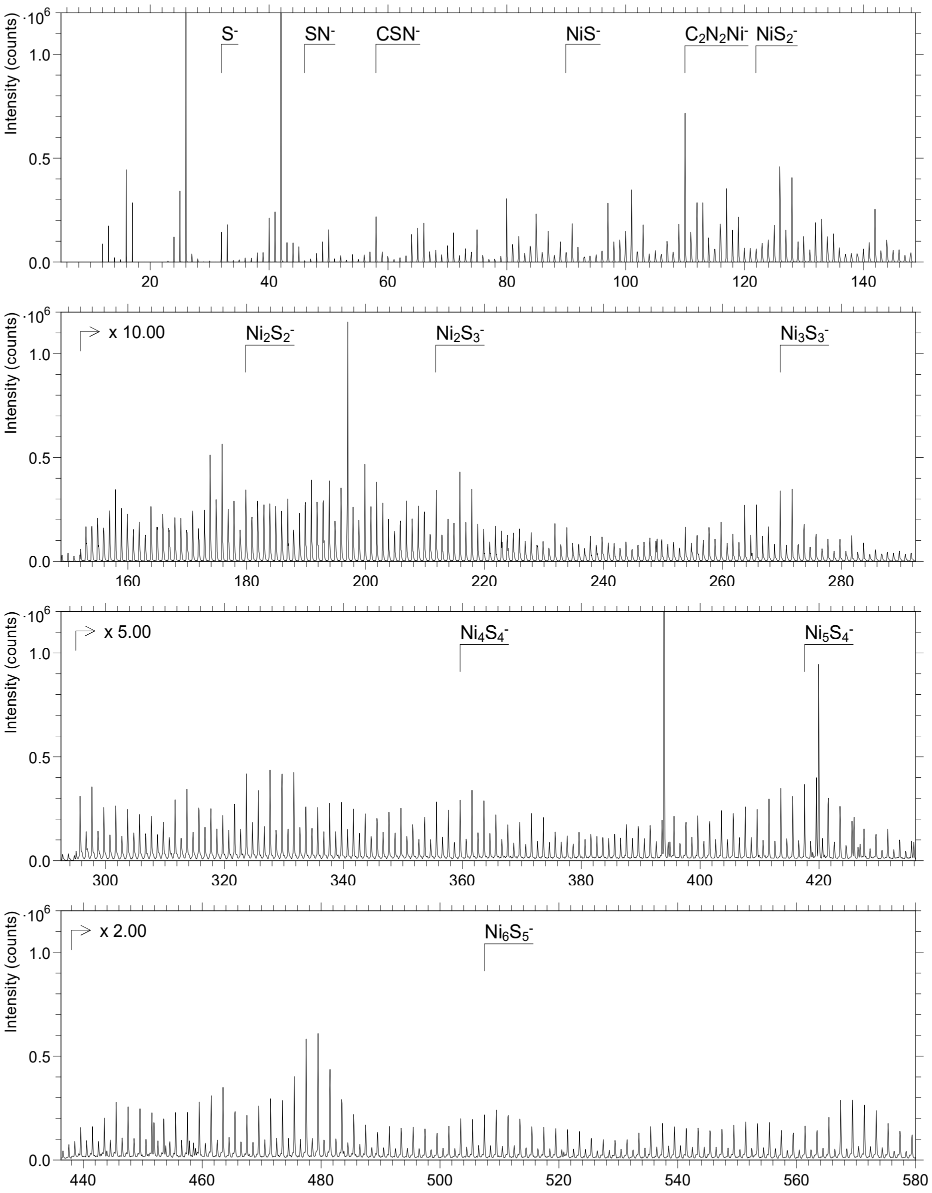

Intensity (counts)
·10^6
S-
SN-
CSN-
NiS-
C2N2Ni-
NiS2-
x 10.00
Ni2S2-
Ni2S3-
Ni3S3-
x 5.00
Ni4S4-
Ni5S4-
x 2.00
Ni6S5-


B

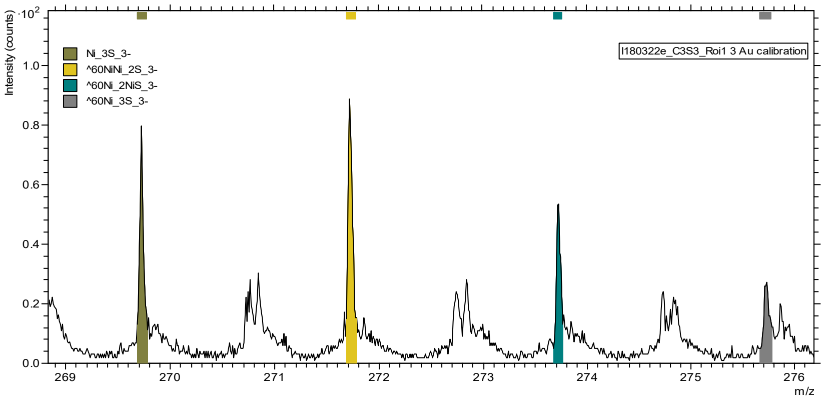

Intensity (counts)
·10^2
Ni_3S_3-
^60NiNi_2S_3-
^60Ni_2NiS_3-
^60Ni_3S_3-
I180322e_C3S3_Roi1 3 Au calibration
m/z

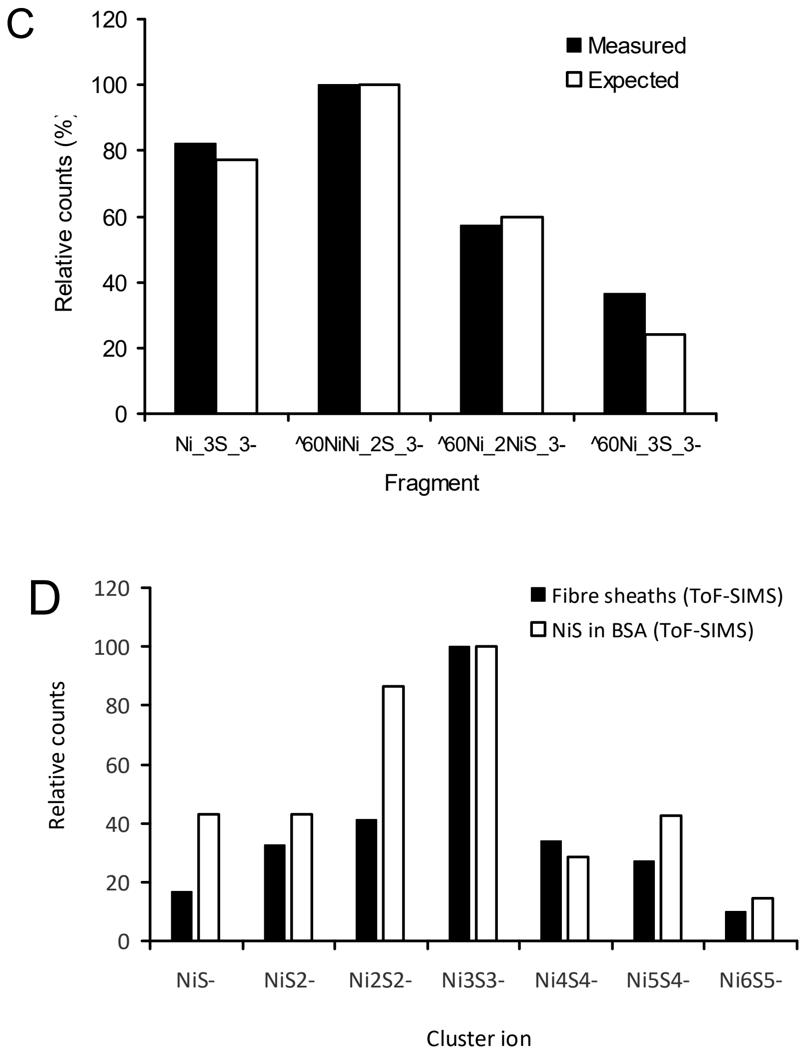


**Supplementary Figure 9.** ToF-SIMS negative mode Ni and S cluster ions from fiber sheaths and a NiS/BSA mixture. A) ToF-SIMS negative mode mass spectrum of a mixture of NiS mineral in BSA. Positions of a selection of important Ni and S cluster ions are indicated. B) Distribution of $Ni_3S_3^-$ isotopomers as a seen in the spectrum of fiber sheaths and C) compared to the expected distribution showing the good fit (similar fits were found for the other $Ni_xS_y$-clusters described, data not shown). D) Distribution of $Ni_xS_y$-cluster ions in fiber sheaths and NiS/BSA mixture (counts were normalized to $Ni_3S_3^-$). Cluster ions with 3 or less Ni atoms were sufficiently mass separated from fragments containing 2 oxygens instead of sulfur such as $Ni_3S_2O_2^-$.

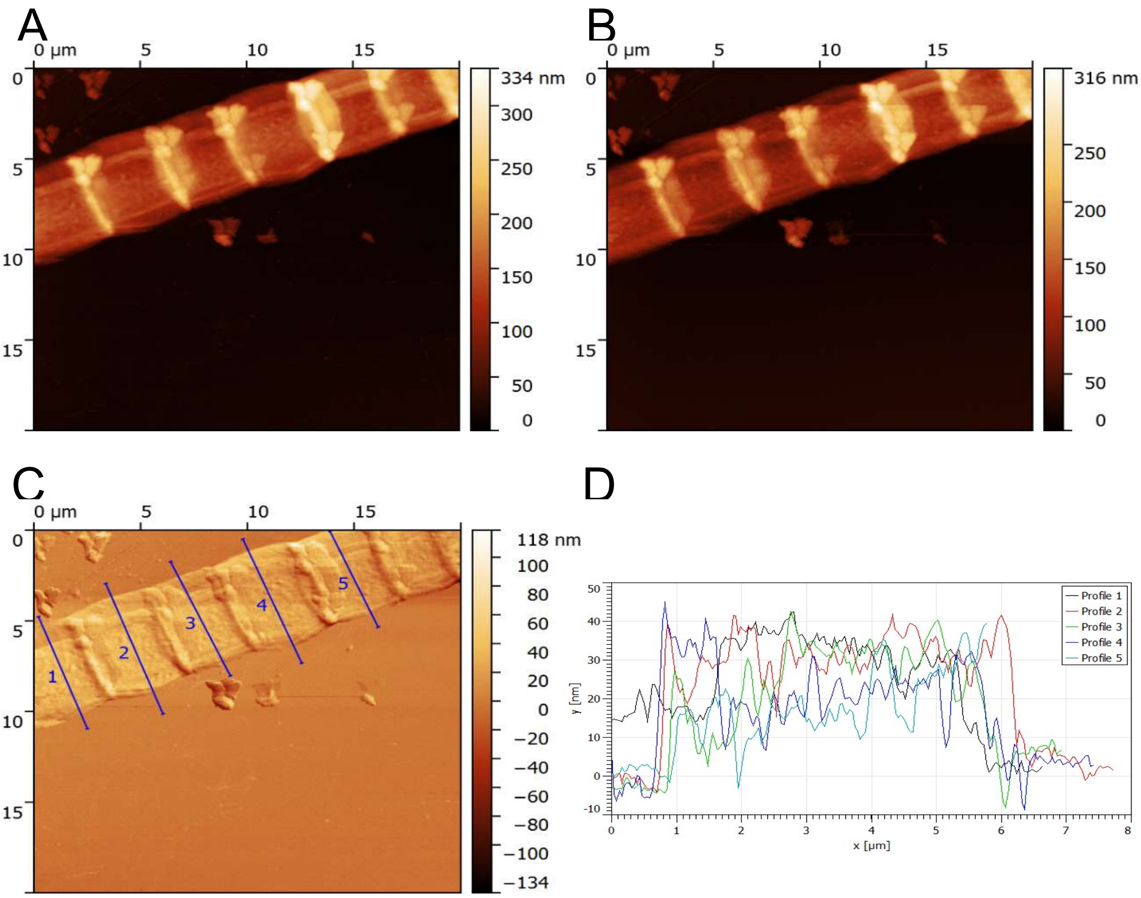


**Supplementary Figure 10.** Example of *in-situ* AFM calibration of ToF-SIMS sputtering depth for fiber sheaths. AFM height map. A) mapping before ToF-SIMS analysis was started. B) mapping after 57 sec of sputtering time (i.e. shortly after the peak in $Ni^+$ counts). C) The differential image between (A) and (B) shows the amount of material removed after 57 sec sputtering. Notice shading of the junctions with higher than average sputtering rates on the side and lower than average sputtering rates on the opposite side. D) Height profiles as indicated by the blue lines in panel (C) showing the lateral variation in sputtering depth. Actual sputtering depths were calculated from height profiles as the average difference in height of the fiber sheath (relative to the wafer surface) before and after sputtering (in this example 22.8 ± 3.2 nm).

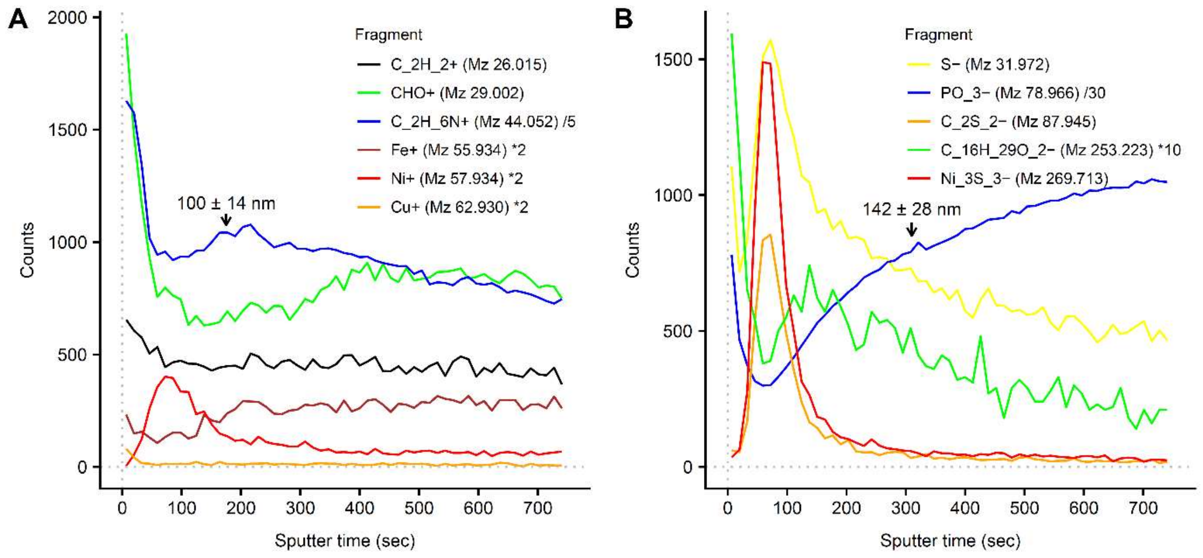


**Supplementary Figure 11.** ToF-SIMS depth profiles of intact cable bacteria demonstrate that the Ni/S group is located in the periplasmic space. Representative ToF-SIMS depth profiles from intact cable bacteria are shown in A) positive mode and B) negative mode. A selection of fragments is shown representing the major biomolecule classes, and counts of individual fragments were scaled to fit all data in a plot. Duplicate runs in both positive and negative modes showed similar profiles. The counts from $Ni_3S_3^-$ are the sum of all $^{58}Ni$ and $^{60}Ni$ isotopologues. The fatty acid fragment ($C_{16}H_{29}O_2^-$) clearly shows two peaks with the first peak at the surface coming most likely from the outer membrane and the broader secondary peak from the cell membrane. Phosphate ($PO_3^-$) also shows a first peak at the surface probably coming from phospholipids and the secondary rise from phospholipids in the cell membrane and poly-phosphate and nucleic acids in the cytoplasm. Arrows indicate sputtering depths as determined with the in-situ AFM in the ToF-SIMS. Average sputtering rate was 0.52 ± 0.05 nm/sec, which places the Ni and S containing peak at approximately 30-40 nm depth in agreement with the expected position of the fibers in intact cable bacteria[2].

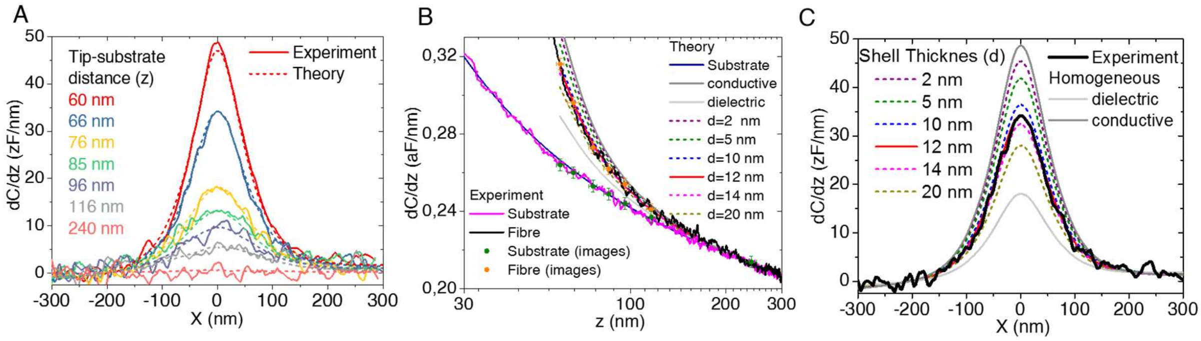


**Supplementary Figure 12.** Additional information on the SDM quantitative analysis shown in Fig. 5 of the main text. A) Capacitance gradient cross-section profiles measured (continuous lines) and calculated (dashed lines) at different tip-substrate distances. The profiles for z=66 nm are those displayed in Fig. 5 of the main text. The parameters used in the calculations are the same as those of Fig. 5 of the main text: H=12.5 μm, W= 3 μm, L=3 μm, l=1 μm, θ=22, R=54 nm, z=60 nm, h=42 nm, w=87 nm, d=12 nm, l=1 μm, $\varepsilon_s = \varepsilon_c = 3$, $\sigma_s = 0$ S/m and $\sigma_c = 2 \cdot 10^3$ S/m. B) Capacitance gradient approach curves measured on a bare part of the substrate (pink continuous line) and on the fiber (black continuous line). The continuous dark blue and red lines correspond to the curves that best fit the experimental results from where the tip geometry (R=54±1 nm, θ=22º±0.5º, $C'_{off}$=112.5±1.5 zF/nm) and shell thickness (d=12±2 nm) have been extracted. The dashed lines represent calculated capacitance gradient approach curves on the fiber for other values of the shell thickness. The continuous dark and grey lines correspond to the homogeneous conductive and dielectric models, respectively. The green and orange symbols correspond to the values measured on the bare substrate and fiber, respectively, on the SDM images at different heights. These values have been used to set the tip-substrate distances of the SDM images. C) Effect of the shell thickness on the calculated capacitance gradient profiles for z=66 nm. The meaning of the lines is the same as in B).

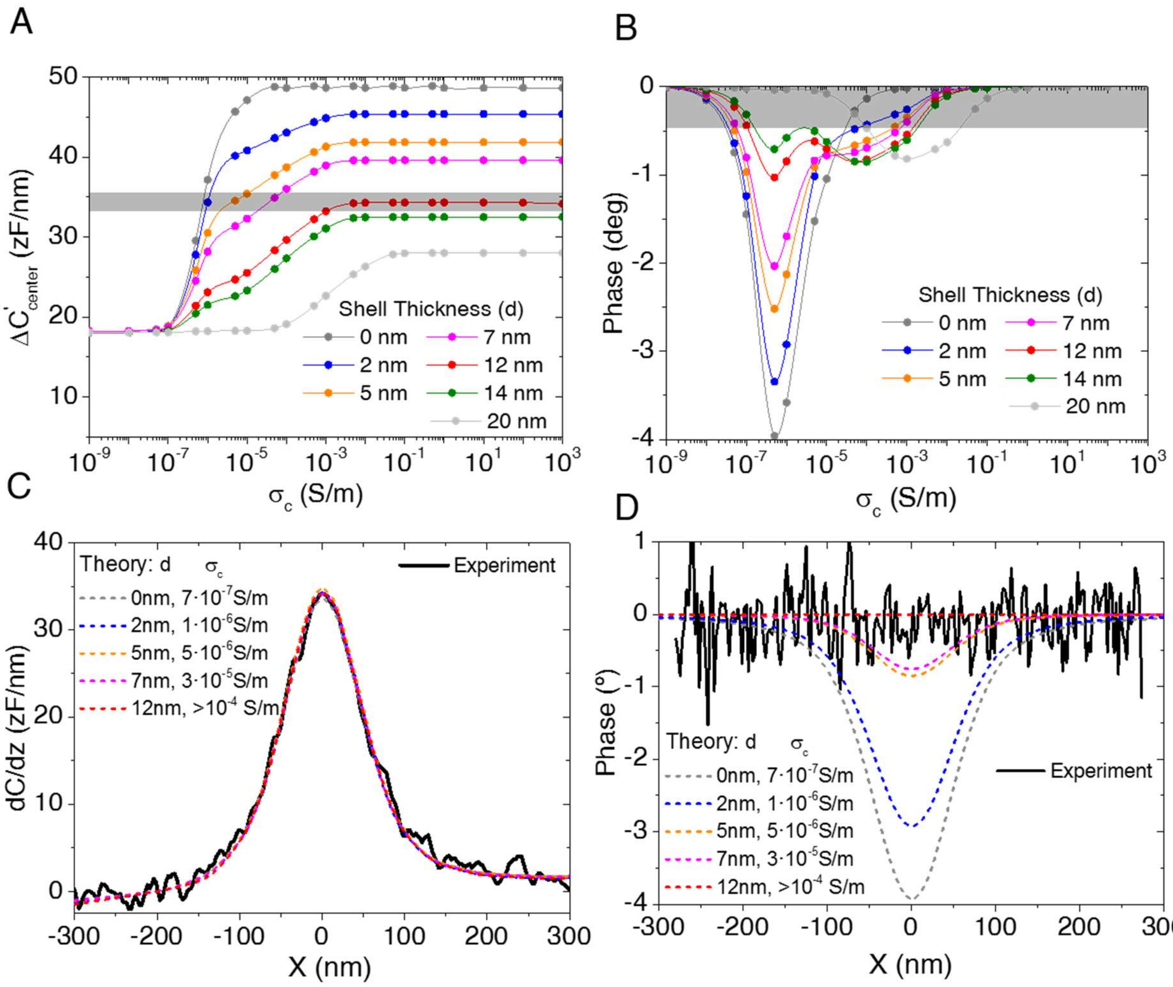


**Supplementary Figure 13.** Additional analysis of the core-shell conductive model used to interpret the SDM measurements shown in Fig. 5 of the main text. A) Amplitude (in capacitance gradient) and B) phase of the electric force contrast calculated for the cylindrical fiber model as a function of the conductivity of the core for different values of the shell thickness. The grey band corresponds to the experimental values extracted from the profiles shown in Fig. 5 of the main text. Calculated capacitance gradient C) and phase D) profiles for the couples of shell thickness-core conductivity that match the experimental measured capacitance gradient contrast in A). The thick lines represent the experimental results (same as in Fig. 5 of the main text). The parameters used in the calculations are the same as those of Fig. 5 of the main text and of Supplementary Figure 12, when not otherwise stated.

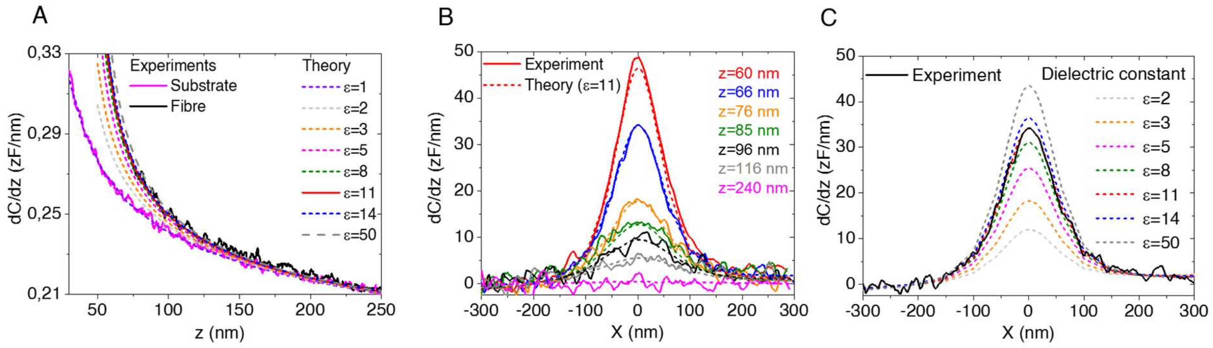


**Supplementary Figure 14.** Analysis of the SDM measurements shown in Fig. 5 of the main text with a homogeneous cylinder dielectric model ($\varepsilon_s = \varepsilon_c = \varepsilon_{eff}$, $\sigma_s = \sigma_c = 0$ S/m). A) Calculated capacitance gradient approach curves on the fiber for different values of the equivalent dielectric constant $\varepsilon_{eff}$ compared to the experimental measured curves on the substrate and fiber (same as in Supplementary Fig 6B, thick lines). The best fit is obtained for $\varepsilon_{eff}=11\pm3$. B) Calculated capacitance gradient profiles for $\varepsilon_{eff}=11$ for different tip-substrate distances, and comparison with the measured profiles (thick lines, same as in Supplementary Figure 6A). C) Calculated capacitance gradient profiles for z=66 nm and different equivalent dielectric constants $\varepsilon_{eff}$, and comparison with the measured profile (thick line, same as in Fig. 5 of the main text). The parameters used in the calculations are the same as those of Fig. 5 of the main text and of Supplementary Figure 6, when not otherwise stated.

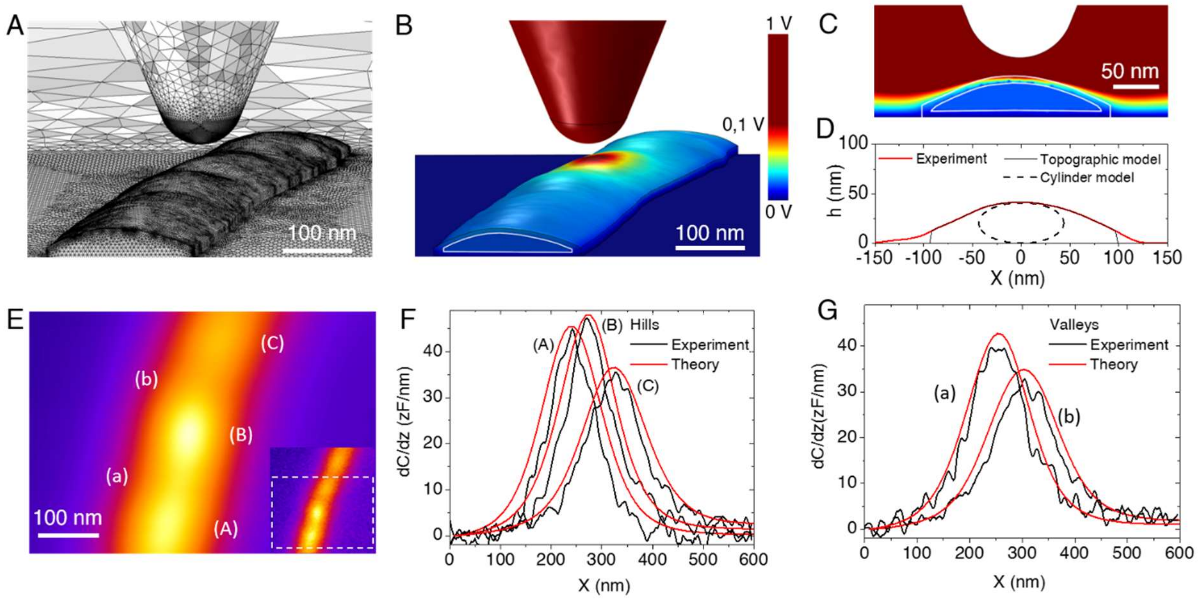


**Supplementary Figure 15.** Analysis of the SDM measurements shown in Fig. 5 of the main text with a geometrical core-shell model reconstructed from the measured topography. A) Detail of the geometry and mesh of the model generated from the measured topography of the fiber (corresponding to the insert in Fig. 5B of the main text). The core-shell structure is defined by assigning a sigmoidal behavior to the dielectric constant and conductivity with plateau representing the shell and core values, respectively. The thickness of the shell corresponds to the center of the sigmoid. B) Example of a calculated electric potential distribution along the fiber for z=66 nm, $\varepsilon_s= \varepsilon_c=3$, $\sigma_s=0$ S/m, $\sigma_c=2\cdot10^3$ S/m and d=10 nm. The tip parameters are the same as those in Fig. 5 of the main text. C) Example of a cross-section electric potential distribution corresponding to B). D) Comparison of the cross-section of the measured topography (red line), the topographically reconstructed fiber model (black continuous line) and of the cylinder model (black dashed line). E) Calculated constant height capacitance gradient SDM image by using the model in A) for the parameters in B). The calculated image corresponds to the area enclosed by the dashed line in the insert. F) and G) Comparison of the calculated capacitance gradient profiles (red lines) with the experimental ones (black lines) on the hills (A,B,C) and valleys (a,b) indicated in E), respectively.

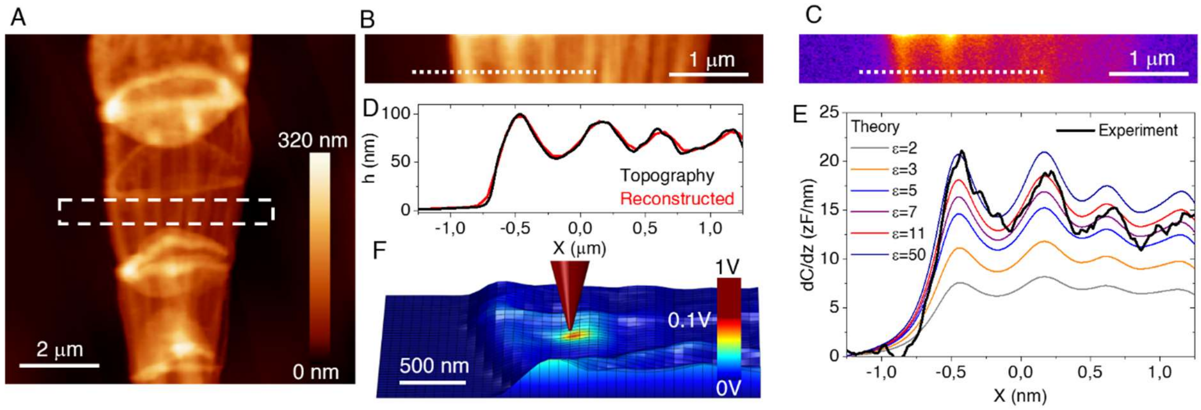


**Supplementary Figure 16.** SDM measurements and analysis on a fiber sheath. A) AFM topographic image of the fiber sheath analyzed. B) and C) AFM topographic and SDM constant height (z=155 nm) images measured on the region enclosed by the dashed rectangle in A). D) and E) Cross-section topographic and capacitance gradient profiles along the dashed lines in B) and C) (thick black lines), respectively. The hills in the image correspond to different fibers. F) Geometrical model reconstructed from the measured topography used in the calculations, with an example of an electric potential distribution overlaid on it. The electrical properties of the fiber sheath have been characterized by an equivalent homogeneous dielectric constant, $\varepsilon_{sheath}$. The tip geometry has been calibrated from a capacitance gradient approach curve on the bare substrate giving R=26±2 nm, θ=22±3°, $C'_{offset}$=109±3 zF/nm. The theoretically predicted capacitance gradient profiles for this model for different values of $\varepsilon_{sheath}$ are shown in E) (thin lines). Most of the hills (fibers) correspond to $\varepsilon_{sheath}$=7−11 in good agreement with the equivalent dielectric constant value found on isolated fibers ($\varepsilon_{eff}$=7−11). For the first hill a higher value is obtained, probably indicating that two fibers are overlaid.

**Supplementary Table 1.** STEM-EDX element compositions for intact cable bacteria and fiber sheaths. Data from two separate imaging sessions are shown (dates in first row).

| | 04-05-17 | | | | 08-08-17 | | | |
|---|---|---|---|---|---|---|---|---|
| | intact cable bacteria N = 3 | | fiber sheath N = 3 | | intact cable bacteria N = 8 | | fiber sheath N = 7 | |
| Elements (atm%) | AVG | SD | AVG | SD | AVG | SD | AVG | SD |
| Carbon | 76.617 | 2.382 | 73.013 | 0.508 | 82.233 | 1.231 | 80.099 | 1.081 |
| Oxygen | 14.370 | 1.185 | 15.880 | 0.322 | 10.854 | 1.394 | 11.319 | 1.205 |
| Nitrogen | 6.133 | 0.250 | 9.283 | 0.224 | 5.411 | 0.547 | 7.333 | 0.477 |
| Phosphorus | 1.120 | 0.607 | 0.663 | 0.100 | 0.833 | 0.167 | 0.736 | 0.150 |
| Sulphur* | 0.510 | 0.035 | 0.350 | 0.225 | 0.483 | 0.078 | 0.329 | 0.023 |
| Zinc | 0.210 | 0.061 | 0.103 | 0.015 | 0.145 | 0.035 | 0.149 | 0.040 |
| Iron | 0.047 | 0.006 | 0.023 | 0.006 | 0.033 | 0.007 | 0.013 | 0.005 |
| Nickel | 0.009 | 0.001 | 0.037 | 0.006 | 0.009 | 0.004 | 0.016 | 0.005 |
| Copper | 0.009 | 0.004 | 0.010 | 0.009 | 0.006 | 0.005 | 0.007 | 0.005 |

*: Sulphur showed interference from a Molybdenum impurity derived most likely from the TEM grid and is therefore uncertain.

**Supplementary Table 2.** Identified fragment ions in positive mode ToF-SIMS of fiber sheaths extracted from cable bacteria. See supplementary Excel file.

**Supplementary Table 3.** Identified fragment ions in negative mode ToF-SIMS of fiber sheaths extracted from cable bacteria. See supplementary Excel file.

**Captions for Movie S1:** 3D tomographic reconstruction of the freeze dried fiber sheath in Fig. 1A

### Supplementary references


1. Burdorf, L. D. W. *et al.* Long-distance electron transport occurs globally in marine sediments. *Biogeosciences* **14**, 683–701 (2017).
2. Cornelissen, R. *et al.* The cell envelope structure of cable bacteria. *Front. Microbiol.* **9**, (2018).
3. Bjerg, J. T. *et al.* Long-distance electron transport in individual, living cable bacteria. *Proc. Natl. Acad. Sci.* **115**, 5786–5791 (2018).
4. Eilers, P. H. C. & Boelens, H. F. M. Baseline correction with asymmetric least squares smoothing. *Leiden University Medical Centre Report*. (2005).
5. Fumagalli, L., Esteban-Ferrer, D., Cuervo, A., Carrascosa, J. L. & Gomila, G. Label-free identification of single dielectric nanoparticles and viruses with ultraweak polarization forces. *Nat. Mater.* **11**, 808–816 (2012).
6. Lozano, H. *et al.* Dielectric constant of flagellin proteins measured by scanning dielectric microscopy. *Nanoscale* **10**, 19188–19194 (2018).
7. Lozano, H., Millán-Solsona, R., Fabregas, R. & Gomila, G. Sizing single nanoscale objects from polarization forces. *Sci. Rep.* **9**, 1–12 (2019).
8. Wang, D. *et al.* Interplay between spherical confinement and particle shape on the self-assembly of rounded cubes. *Nat. Commun.* **9**, 2228 (2018).
9. van Aarle, W. *et al.* The ASTRA toolbox: A platform for advanced algorithm development in electron tomography. *Ultramicroscopy* **157**, 35–47 (2015).
10. Gianoncelli, A., Kourousias, G., Merolle, L., Altissimo, M. & Bianco, A. Current status of the TwinMic beamline at Elettra: A soft X-ray transmission and emission microscopy station. *J. Synchrotron Radiat.* **23**, 1526–1537 (2016).

11. Solé, V. A., Papillon, E., Cotte, M., Walter, Ph. & Susini, J. A multiplatform code for the analysis of energy-dispersive X-ray fluorescence spectra. *Spectrochim. Acta Part B At. Spectrosc.* **62**, 63–68 (2007).
12. Polerecky, L. *et al.* Look@NanoSIMS - a tool for the analysis of nanoSIMS data in environmental microbiology. *Environ. Microbiol.* **14**, 1009–1023 (2012).
13. Meysman, F. J. R. *et al.* A highly conductive fibre network enables centimetre-scale electron transport in multicellular cable bacteria. *Nat. Commun.* **10**, 1–8 (2019).
14. Baugh, L. *et al.* Probing the orientation of surface-immobilized protein G B1 using ToF-SIMS, sum frequency generation, and NEXAFS spectroscopy. *Langmuir* **26**, 16434–16441 (2010).
15. Goacher, R. E., Jeremic, D. & Master, E. R. Expanding the library of secondary ions that distinguish lignin and polysaccharides in time-of-flight secondary ion mass spectrometry analysis of wood. *Anal. Chem.* **83**, 804–812 (2011).
16. Wei, W. *et al.* Characterization of syntrophic Geobacter communities using ToF-SIMS. *Biointerphases* **12**, 05G601 (2017).
17. Lebec, V., Boujday, S., Poleunis, C., Pradier, C.-M. & Delcorte, A. Time-of-flight secondary ion mass spectrometry investigation of the orientation of adsorbed antibodies on SAMs correlated to biorecognition tests. *J. Phys. Chem. C* **118**, 2085–2092 (2014).
18. Ding, Y. *et al.* In Situ Molecular Imaging of the Biofilm and Its Matrix. *Anal. Chem.* **88**, 11244–11252 (2016).
19. Lee, C.-Y., Harbers, G. M., Grainger, D. W., Gamble, L. J. & Castner, D. G. Fluorescence, XPS, and TOF-SIMS Surface Chemical State Image Analysis of DNA Microarrays. *J. Am. Chem. Soc.* **129**, 9429–9438 (2007).

20. May, C. J., Canavan, H. E. & Castner, D. G. Quantitative X-ray Photoelectron Spectroscopy and Time-of-Flight Secondary Ion Mass Spectrometry Characterization of the Components in DNA. *Anal. Chem.* **76**, 1114–1122 (2004).
21. Lovering, A. L., Safadi, S. S. & Strynadka, N. C. J. Structural perspective of peptidoglycan biosynthesis and assembly. *Annu. Rev. Biochem.* **81**, 451–478 (2012).
22. Geerlings, N. M. J., Zetsche, E.-M., Hidalgo Martinez, S., Middelburg, J. J. & Meysman, F. J. R. Mineral formation induced by cable bacteria performing long-distance electron transport in marine sediments. *Biogeosciences Discuss.* 1–35 (2018) doi:10.5194/bg-2018-444.
23. Franquet, A. *et al.* Self focusing SIMS: Probing thin film composition in very confined volumes. *Appl. Surf. Sci.* **365**, 143–152 (2016).
24. Feld, H., Rading, D., Leute, A. & Benninghoven, A. Comparative investigations of the secondary ion emission of metal complexes under MeV and keV ion bombardment. *Org. Mass Spectrom.* **28**, 841–852 (1993).
25. El Nakat, J. H., Dance, I. G., Fisher, K. J., Rice, D. & Willett, G. D. Laser-ablation FTICR (Fourier-transform ICR) mass spectrometry of metal sulfides: gaseous anionic nickel-sulfur [NixSy] clusters. *J. Am. Chem. Soc.* **113**, 5141–5148 (1991).
26. Jones, T. B. *Electromechanics of particles*. (Cambridge University Press, 1995).
27. Cuervo, A. *et al.* Direct measurement of the dielectric polarization properties of DNA. *Proc. Natl. Acad. Sci.* **111**, E3624–E3630 (2014).
28. Dols-Perez, A., Gramse, G., Calò, A., Gomila, G. & Fumagalli, L. Nanoscale electric polarizability of ultrathin biolayers on insulating substrates by electrostatic force microscopy. *Nanoscale* **7**, 18327–18336 (2015).
29. Checa, M. *et al.* Mapping the dielectric constant of a single bacterial cell at the nanoscale with scanning dielectric force volume microscopy. *Nanoscale* **11**, 20809–20819 (2019).

30. Fabregas, R. & Gomila, G. Dielectric nanotomography based on electrostatic force microscopy: A numerical analysis. *J. Appl. Phys.* **127**, 024301 (2020).
